# Fork Reversal Safeguards Epigenetic Inheritance during DNA Replication Under Stress

**DOI:** 10.1101/2025.09.21.677631

**Authors:** Qiong Wu, Caixian Zhou, Yue Dou, Nadia I. Martin, Maria Gwit, Tae Hee Lee, Mark Hedglin, Roséa Chen, Ke Zhu, Tianpeng Zhang, Shangming Tang, Tian Zhang, Tyler M. Weaver, Wenpeng Liu

**Author notes:** These authors contributed equally (Qiong Wu, Caixian Zhou). Correspondence: W. L.

## Abstract

During DNA replication, epigenetic information carried by histone modifications is faithfully propagated and re-established on sister chromatids, ensuring cell identity. Chromatin reassembly is tightly coupled to DNA replication, therefore whether and how perturbations to DNA replication affects the fidelity of epigenetic inheritance remain poorly understood. In this study, we reveal a critical role for replication fork reversal in maintaining the transmission of epigenetic information under replication stress. We identify that cells defective in fork reversal exhibit reduced nucleosome density at replication forks, accompanied by the loss of parental histones during their transfer onto nascent DNA. Mechanistically, we demonstrate that PrimPol activation leads to single-stranded DNA gaps in fork reversal deficient cells, and that subsequent PARylation (poly ADP-ribosylation) and DNA–protein crosslinking on these gaps evicts nucleosomes. Our findings demonstrate that replication fork reversal, a widespread physiological process, is not only essential for preserving genome integrity but also for safeguarding epigenetic stability.

## Introduction

In eukaryotic cells, DNA replication is not merely the duplication of genetic material; it is also a critical process for the faithful transmission of epigenetic information^1^. During each S phase of the cell cycle, DNA must be precisely duplicated, and chromatin structure must be accurately reassembled to ensure cell identity, proper gene expression, and overall genome stability^2,3^. This chromatin-based inheritance, referred to as epigenetics, encompasses histone modifications, DNA methylation, and regulation by non-coding RNAs^4^. Programmed regulation of epigenetic states is essential for cell differentiation and organismal development^5^, whereas aberrant epigenetic changes can drive diseases and tumorigenesis^6–10^. In cancer therapy, epigenetic exacerbates tumor cell heterogeneity and promotes drug resistance^11^. Therefore, understanding how epigenetic information is transferred during DNA replication is of fundamental importance.

During DNA replication, nucleosomes ahead of the fork are disassembled by the histone chaperone FACT (FAcilitates Chromatin Transcription) complex into one hexamer, which contains a H3–H4 tetramer and a H2A–H2B dimer. Next, the hexamer is recycled to its original positions on one of the sister chromatid by multiple histone chaperones ^1,12–20^. With the assistance of histone chaperones, like ASF1 and CAF1, recycled parental histones are incorporated together with newly synthesized histones to assemble nucleosomes on nascent DNA^1,21–32^. Post-translational modifications (PTMs) on parental histones serve as templates that are copied onto adjacent new histones through a “read–write” mechanism, thereby completing the replication of epigenetic information^1,33–37^. Replisome proteins such as MCM2, PCNA, DNA polymerases ε/α, Tof1, Mrc1, Csm3, and Ctf4 contribute to parental histone transfer^25–27,35,38–48^. As a result, DNA replication exerts a profound influence on epigenetic stability. For example, depletion of Pol E3/E4, subunits of leading-strand histone chaperone polymerase ε, disrupt H3K9me3 marked parental histones recycling, thereby activating expression of LINE-1 (Long Interspersed Element-1) retrotransposon^49^. Conversely, mutations in lagging-strand histone chaperones, such as MCM2-2A, lead to parental histone accumulation on the leading strand^38,39^. Simultaneous disruption of both leading- and lagging-strand histone chaperones results in the loss of parental histones and histone PTMs, impairing stem cell differentiation and viability^50^.

DNA replication frequently encounters obstacles such as transcription–replication conflicts and repetitive sequences^3,51^. Replication stress response pathways are activated to mitigate the replication obstacles^3,51^. Among these pathways, the generation of single-stranded DNA (ssDNA) and fork reversal are the two most common structural intermediates, occurring in diverse cell types in response to various forms of replication stress^52,53^. We and others previously identified that ssDNA can be processed by RAD51-mediated strand invasion together with multiple fork remodelers, including SMARCAL1, ZRANB3, and HLTF, to form reversed forks^53–64^. This process stabilizes replication intermediates and prevents excessive ssDNA accumulation to promote proper DNA replication. When fork reversal is impaired, cells activate PrimPol to reinitiate DNA synthesis downstream of the leading strand in a process known as repriming^65,66^. Although replication can resume, this generates abundant ssDNA gaps on nascent DNA that pose a threat to chromatin structure and genome stability^67–69^.

Under stressed conditions, DNA replication progress slowly compared to non-stressed conditions, rather than completely stop^70^. It is unclear how fork reversal or ssDNA gaps influence parental histone recycling and nucleosome assembly. In this study, we demonstrate that replication fork reversal promotes parental histone recycling and nucleosome assembly on sister chromatids by suppressing ssDNA formation, thereby preserving epigenetic stability. In cells lacking the ability to undergo fork reversal, activation of the PrimPol primase leads to repriming and the generation of ssDNA gaps, which ultimately impairs recycling of parental histones and histone PTMs. We further show that impaired histone recycling results from the accumulation of ssDNA that triggers PARP1-mediated PARylation and DNA–protein crosslink formation, both of which inhibit nucleosome assembly. Together, our findings reveal a critical role for replication fork reversal in safeguarding epigenetic stability under replication stress.

## Results

### Replication fork reversal deficiency reduces nucleosome density on nascent DNA under replication stress

To investigate the impact of replication fork reversal on replication-associated nucleosome assembly at replication forks, we performed SILAC-iPOND-MS (Stable Isotope Labeling by Amino acids in Cell Culture - Isolation of Proteins on Nascent DNA - Mass Spectrometry) analysis in wild-type (WT) U2OS cells and U2OS cells lacking the three major fork reversal enzymes, including SMARCAL1, HLTF, ZRANB3 (termed as FRKO) to examine protein composition on nascent DNA, upon hydroxyurea (HU)-induced replication stress (Figure 1A). The SILAC-iPOND-MS analysis identified that FRKO cells have a markedly decreased abundance of nearly all histones on nascent DNA compared to WT cells (Figure 1A). Notably, the levels of the MCM2–7 helicase complex remained unchanged between WT and FRKO cells, indicating that histone reduction was not due to a decrease in replication fork number (Figure 1A). In addition, compared to WT cells, FRKO cells showed mild increased levels of EdU (5-Ethynyl-2’-deoxyuridine) incorporation (Figure S1A), indicating that histone reduction in FRKO cells is not the result of reduced replication progression. Immunofluorescence analysis of total chromatin bound histone H3 showed no difference between fork reversal–deficient and wild-type cells (Figure S1B–E), identifying that histone loss was restricted to nascent DNA.

**Figure 1.**
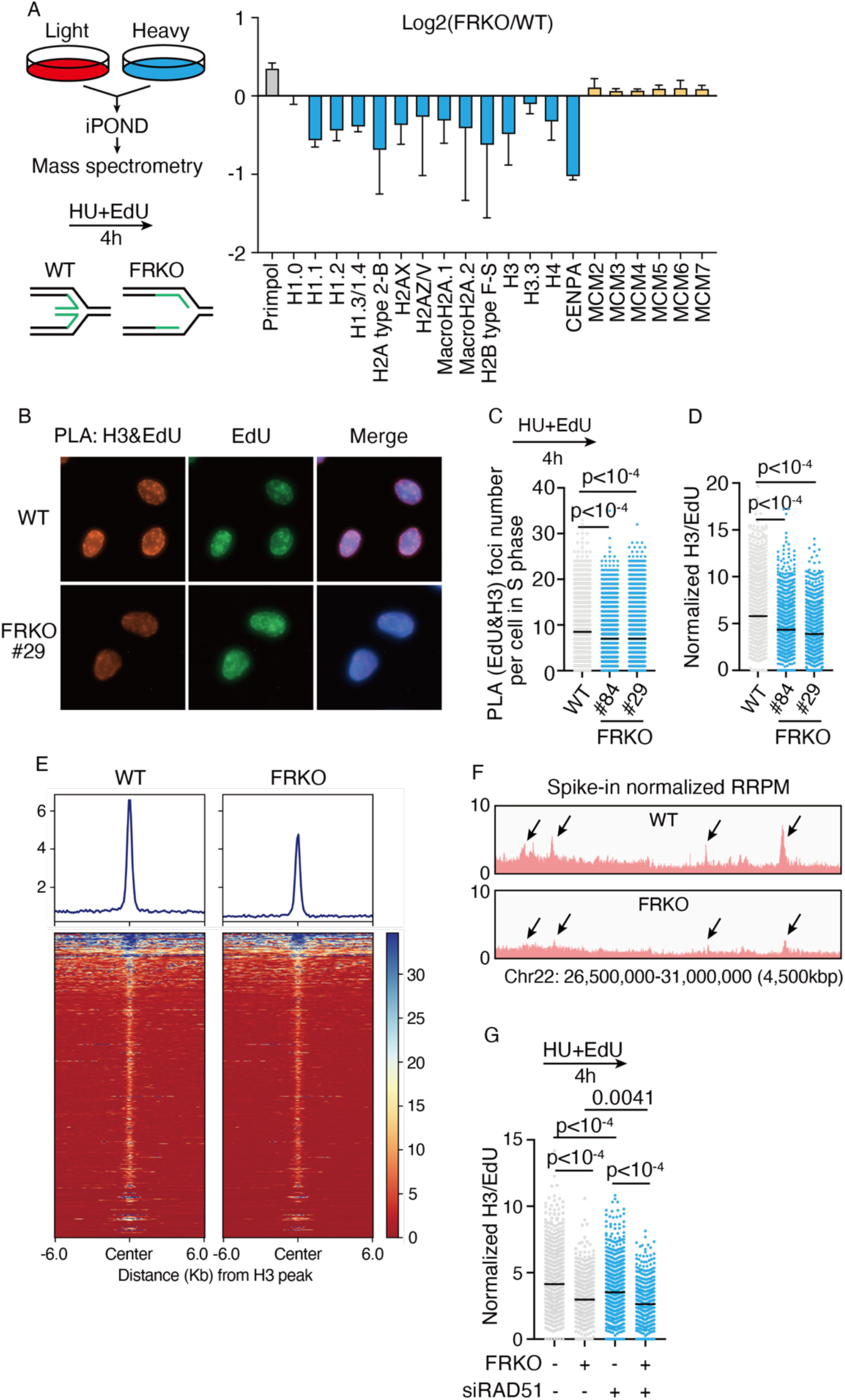
Fork reversal deficiency reduces nucleosome density on nascent DNA. A) iPOND-SILAC-MS analysis of proteins bound to nascent DNA in HU-treated wild-type (WT) and fork reversal enzyme knockout (FRKO) U2OS cells. WT and FRKO cells were labeled with light and heavy isotopes for ≥4 weeks, pulse-labeled with EdU and treated with 1mM HU for 4 h prior to iPOND. N = 4 biological replicates; error bars represent SD. B–D) PLA for EdU and H3 in WT or FRKO cells labeled with 10 μM EdU and treated with 1 mM HU. B) Representative images (Texas Red represents PLA signal; green represents EdU signal). C) PLA foci number per cell; each dot represents PLA foci number from a single cell. D) PLA signal normalized to EdU intensity. ≥1,000 cells/sample were analyzed with MetaXpress software. N ≥ 10. p-values from Kruskal–Wallis test. E) Heatmaps of H3 ChOR-seq signal in WT and FRKO cells treated with HU, normalized to spike-in chromatin from D. melanogaster S2 cells. N = 2. F) Genome browser views of H3 ChOR-seq from the same experiment in (E), normalized to reads per million using exogenous spike-in. N = 2. G) PLA for EdU and H3 in WT or FRKO cells with siRAD51 knockdown. N = 3. p-values from Kruskal–Wallis test.

To validate the mass spectrometry results, we pulse labeled nascent DNA with EdU, induced replication stress with HU, and quantified histone H3 on nascent DNA using a proximity ligation assay (PLA)^71^ in WT and two clones of FRKO U2OS cells. Since FRKO cells incorporate EdU more rapidly than wild-type cells (Figure S1A), direct comparison of PLA signals would not accurately reflect histone abundance on nascent DNA (Figure 1B, 1C). Therefore, we normalized PLA signals to EdU intensity, yielding a value representing histone abundance per unit length of replicated nascent DNA (Figure 1D). Using this approach, we observed a significant decrease in H3 levels on nascent DNA in two FRKO clones. Additional PLA analyses identified a marked decrease in H2A, H2B, and H4 abundance, consistent with the observations for H3 (Figure 1B-D), indicating the FRKO cells have a decreased abundance of all four core histones rather than loss of individual histone types (Figure S1F-H). To further validate these results, we used ChOR (Chromatin Occupancy after Replication)-seq, a high-throughput sequencing technique to profiling the occupancy of histone (or modifications) and chromatin-associated proteins on newly replicated DNA^72^. The ChOR-seq analysis also identified a significant reduction in histone H3 abundance on nascent DNA in the FRKO cells (Figure 1E-F, Figure S1I-J). Together, the PLA and ChOR-seq analysis is consistent with the replication fork proteomics experiments that histone abundance is reduced, suggesting low nucleosome density on nascent DNA in FRKO cells (Figure 1A).

These experiments were performed in U2OS cells with co-depletion of SMARCAL1, ZRANB3, and HLTF. To determine which enzyme loss dominates the phenotype, we performed PLA analysis in two clones for every single knockout of SMARCAL1, ZRANB3, and HLTF in U2OS cells (Figure S2A–C). Notably, among these single knockout clones, both clones for HLTF knockout consistently show significantly lower H3 level on nascent DNA (Figure S2A and S2C), suggesting HLTF depletion dominates the phenotype in FRKO U2OS cells. Additionally, SMARCAL1 and ZRANB3 single KO also shows a mild decrease in H3 level on nascent DNA (Figure S2A-C). FRKO summarizes all three single depletions (Figure S2C). Consistent with observations in fork reversal enzyme knockout cells, knockdown of RAD51, which is also an essential factor for fork reversal, led to low nucleosome density on nascent DNA (Figure 1G and Figure S2D).

Notably, PLA and ChOR-seq showed no reduction in H3 levels in FRKO cells without treatment with HU (Figure S2E-G), supporting that histone loss arises as a result of replication stress. Together, these data indicate that fork reversal machinery maintains nucleosome density on nascent DNA during conditions of replication stress (Figure S2H).

### Replication fork reversal deficiency leads to parental histone loss

During DNA replication, parental histones positioned ahead of the fork are disassembled and subsequently recycled onto daughter DNA through highly coordinated mechanisms, where they combine with newly synthesized histones to form nucleosomes. Under conditions of persistent replication stress, the reduced nucleosome density observed on nascent DNA in FRKO cells may arise from either impaired parental histone recycling or defective incorporation of new histones, both of which threaten faithful epigenetic inheritance. Importantly, the deposition of certain histone variants, such as CENP-A, macroH2A, and H3.3, occurs independently of DNA replication ^73–75^. Thus, detection of these variants on nascent DNA reflects inheritance of parental histones, and their loss can be attributed to defective parental histone recycling (Figure 1A).

Prior work identified that specific histone post-translational modifications (PTMs), such as di- and trimethylation of H3K9, H3K79, and H4K20, are established on new histones many hours after replication or during the subsequent cell cycle, making them markers of parental histones during replication^31,39,49,50,76–78^. To assess whether parental histone recycling was impaired in FRKO cells under replication stress, we combined iPOND-MS with chemical derivatization of histone treatment to examine histone modifications at replication forks (Figure 2A)^79^. Upon HU-induced replication stress, parental histone PTMs such as H4K20me2, H3K79me2, H3K27me3, and H3K9me3 were markedly reduced on nascent DNA in fork reversal–deficient cells (Figure 2B, Figure S3A-D). In contrast, histone PTMs characteristic of newly synthesized histone modifications^49,77^, such as H4K12ac, H4K5ac, H3K56ac, H3K9ac were either increased or remained unchanged (Figure 2B, Figure S3E-H). To validate H3K9me3 reduction in iPOND-MS results, we performed PLA assays to examine H3K9me3 levels on nascent DNA. We found that H3K9me3 levels on nascent DNA are reduced, consistent with iPOND-MS results (Figure 2C).

**Figure 2.**
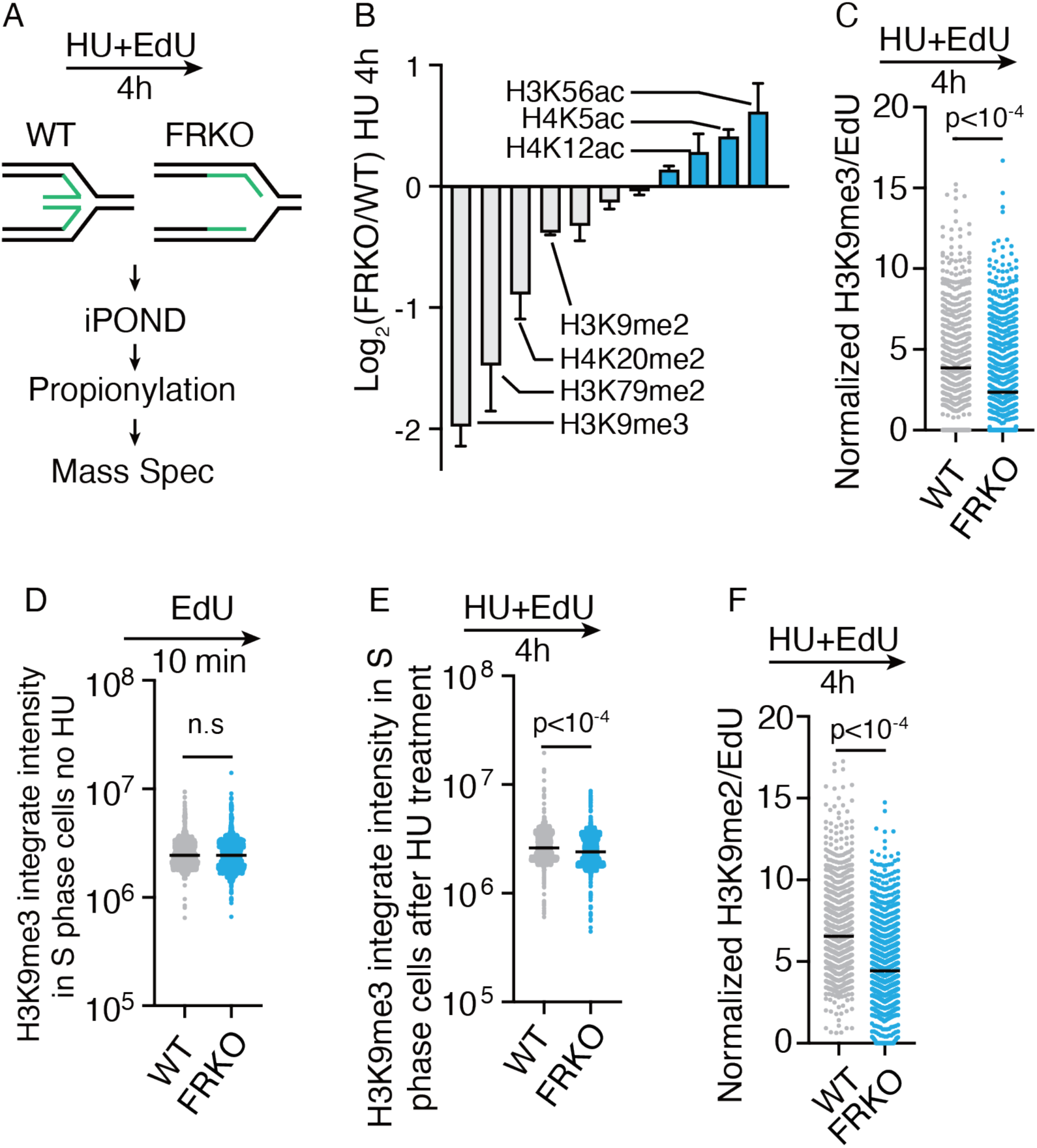
Fork reversal deficiency leads to parental histone loss on nascent DNA. A) Schematic of iPOND–propionylation-MS workflow to detect histone modifications on nascent DNA in WT and FRKO U2OS cells treated with EdU together with 1mM HU for 4 h. B) Log₂ fold-change of histone modifications in FRKO vs. WT cells. Modification-specific peptide counts were normalized to unique peptide numbers, and then FRKO/WT ratios plotted on log₂ scale. N = 3. C) PLA for EdU and H3K9me3 in WT and FRKO cells treated as in (A). PLA signals normalized to EdU intensity. ≥1,000 cells/sample. N = 3. p-values from two-tailed unpaired t-test. D–E) Immunofluorescence for H3K9me3 in S-phase WT and FRKO cells. D) Cells pulse-labeled with EdU (10 min). E) Cells labeled with EdU and treated with HU (4 h). Quantification by QIBC. N = 3. p-values from two-tailed unpaired t-test. F) PLA for EdU and H3K9me2 in WT and FRKO cells treated as in (A). PLA signals normalized to EdU intensity. ≥1,000 cells/sample. N = 3. p-values from two-tailed unpaired t-test.

H3K9me3 is a parental histone PTMs that are associated with constitutive heterochromatin, which is lost upon replication stress in FRKO cells. To distinguish whether the reduction of H3K9me3 arises from replication stress or from preexisting global changes in FRKO cells, we performed immunofluorescence analysis. A slight decrease in cellular H3K9me3 level was observed only in S phase FRKO cells following HU treatment, rather than unperturbed condition (Figure 2D–E), indicating that the loss of this repressive mark in fork reversal–deficient cells is specifically attributable to replication stress. These results indicate the loss of parental histone during DNA replication in FRKO cells under stressed conditions.

H3K9me3 and H3K27me3 are parental histone–specific PTMs that also indicate compact chromatin. We also detected a decrease in the abundance of another compaction marker, H3K9me2 on nascent DNA in fork reversal–deficient cells (Figure 2B, 2F, Figure S3I), while the cellular H3K9me2 levels decrease also depends on HU treatment (Figure S3J-K). These findings are consistent with our model in which nucleosome density is reduced on nascent DNA in FRKO cells upon replication stress.

### PrimPol-mediated repriming causes nucleosome density reduction

Histone chaperones are responsible for the deposition of newly synthesized histones and the recycling of parental histones behind the replication fork. Impaired recruitment of histone chaperones to the replication fork could explain the robust decrease in nucleosome density and parental histone recycling observed in FRKO cells during replication stress. Analysis of histone chaperone localization from our SILAC-iPOND-MS analysis revealed that almost all known histone chaperones properly localize to the replication fork in both WT and FRKO cells (Figure 3A), suggesting that the loss of parental histones and reduced nucleosome density are not due to defects of histone chaperone recruitment.

**Figure 3.**
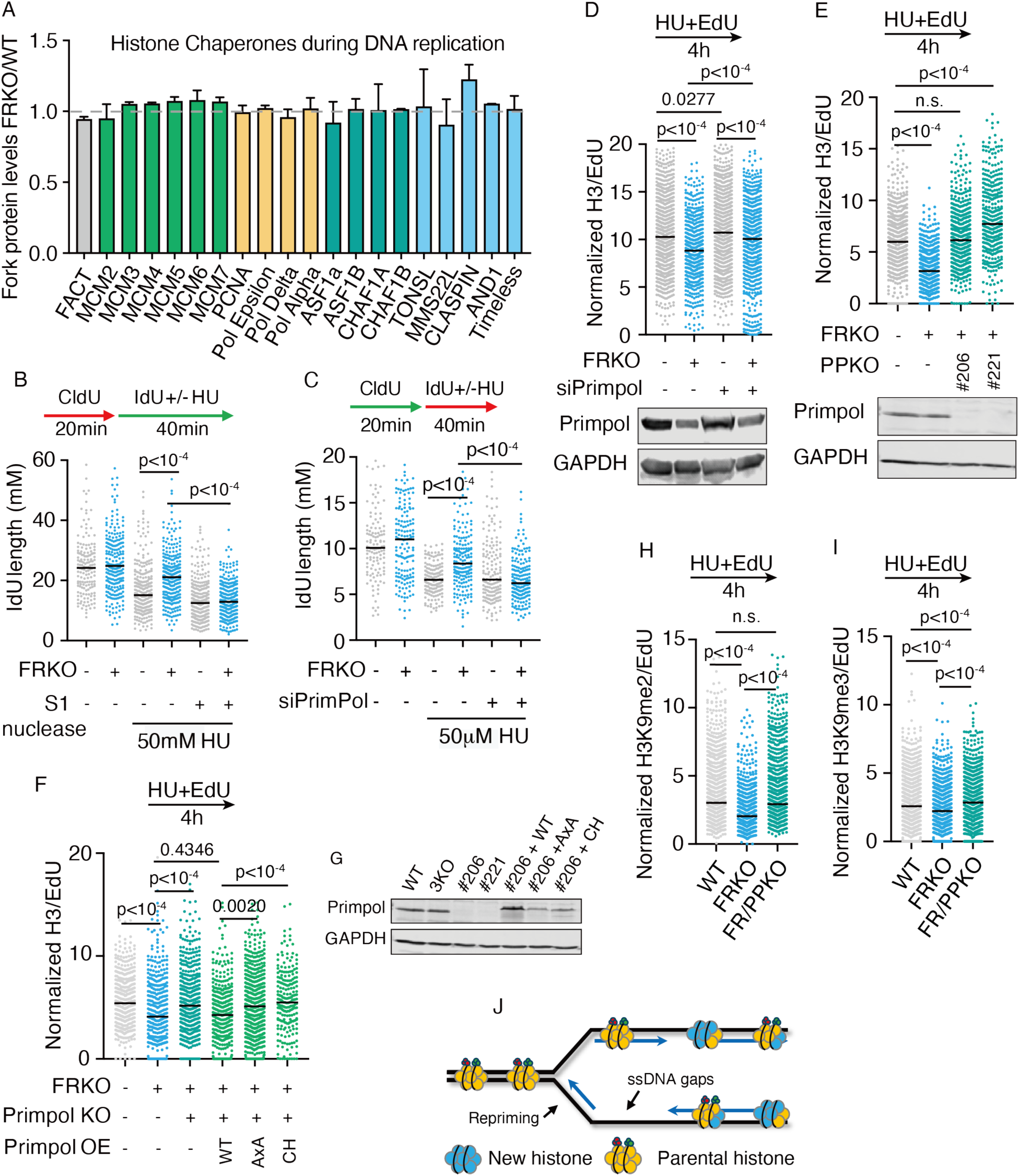
PrimPol deletion rescues low nucleosome density. A) iPOND-SILAC-MS analysis of histone chaperones on nascent DNA (data from Figure 1A). N = 4, error bars generated by Standard deviation. B-C) S1 nuclease-treated DNA fiber assay to detect ssDNA gaps. N = 3. p-values from Kruskal–Wallis test. D-F) PLA for EdU and histone H3 in indicated conditions. PLA signals normalized to EdU intensity; ≥1,000 cells/sample. N ≥ 3. p-values from Kruskal–Wallis test. G) Western blot for PrimPol in KO and complemented cells. H-I) PLA for EdU and histone modifications in indicated conditions. H) H3K9me2; I) H3K9me3. PLA signals normalized to EdU intensity; ≥1,000 cells/sample. N ≥ 3. p-values from Kruskal–Wallis test. J) Model schematic.

We noted that PrimPol (Primase And DNA Directed Polymerase) levels at replication forks were elevated after HU treatment in FRKO cells (Figure 1A), consistent with previous reports of PrimPol activation under fork reversal defects^65,66,80^. We hypothesized that PrimPol activation may contribute to loss of nucleosome density. Following HU treatment, FRKO cells exhibited elevated ssDNA gaps on nascent DNA (Figure 3B). PrimPol depletion can rescue the ssDNA gaps, suggesting PrimPol repriming is activated in FRKO cells and these gaps are PrimPol dependent (Figure 3C). To test whether PrimPol activation was responsible for reduced nucleosome density under replication stress, we depleted PrimPol in FRKO cells by siRNA or CRISPR-Cas9 and measured nucleosome density on nascent DNA by PLA assay (Figure 3D-E). PrimPol depletion in both conditions rescued low nucleosome density in FRKO cells. To determine if the reduction in nucleosome density requires PrimPol catalytic activities, we reintroduced either wild-type PrimPol or a catalytically inactive point mutant (AxA (catalytic inactive), CH (primase dead, polymerase competent) mutants) into FR/PrimPol-knockout (FR/PPKO) cells. Only wild-type PrimPol was sufficient to restore nucleosome density, demonstrating that PrimPol repriming activity drives the loss of nucleosome density (Figure 3F-G).

Next, to determine if the parental-specific histone modification loss in FRKO cells results from PrimPol repriming, we performed iPOND-MS for histone modifications in FR/PPKO cells (Figure S4A-D). Our results demonstrate that PrimPol depletion restores H4K20me2 and H3K79me2 levels (Figure S4A-B) and reverses the increase of H4K5Ac and H3K56Ac in FRKO cells (Figure S4C-D). While we did not detect H3K9me2/3 peptides in iPOND-MS, PLA assays for H3K9me2/3 on nascent DNA confirmed restoration of H3K9me2 and H3K9me3 levels (Figure 3H-I).

Collectively, these results demonstrate that when fork reversal is absent, PrimPol-mediated repriming leads to loss of parental histones and a reduction in nucleosome density on nascent DNA (Figure 3J).

### Single-stranded DNA gaps cause low nucleosome density in cells

To determine if the low nucleosome density attribute to the elevated levels of ssDNA gaps that mediated by PrimPol, or enriched PrimPol levels per se, we examined nucleosome density in other previously defined ssDNA gaps contexts, such as BRCA1 or BRCA2 deficient cells. BRCA1 and BRCA2 are key fork protection factors, and BRCA1- or BRCA2-deficiency cells combined with HU treatment leads to nucleolytic degradation of nascent DNA and ssDNA gap formation; PrimPol repriming is also activated in the absence of BRCA2^67,81–85^. We performed PLA assay for EdU and H3 in BRCA1 deficient UWB1.289 cells, and UWB1.289 cells complement with wild-type BRCA1. Compared to complementation cells, we observed low nucleosome density in UWB1.289 cells (Figure 4A, Figure S5A). Similarly, we also observed low nucleosome density in other BRCA1- or BRCA2-deficient cells lines (Figure 4B–D, Figure S5B–D). These findings are consistent with prior work showing that histone H3 levels are decreased on nascent DNA in BRCA2 knockout HeLa cells^85^.

**Figure 4.**
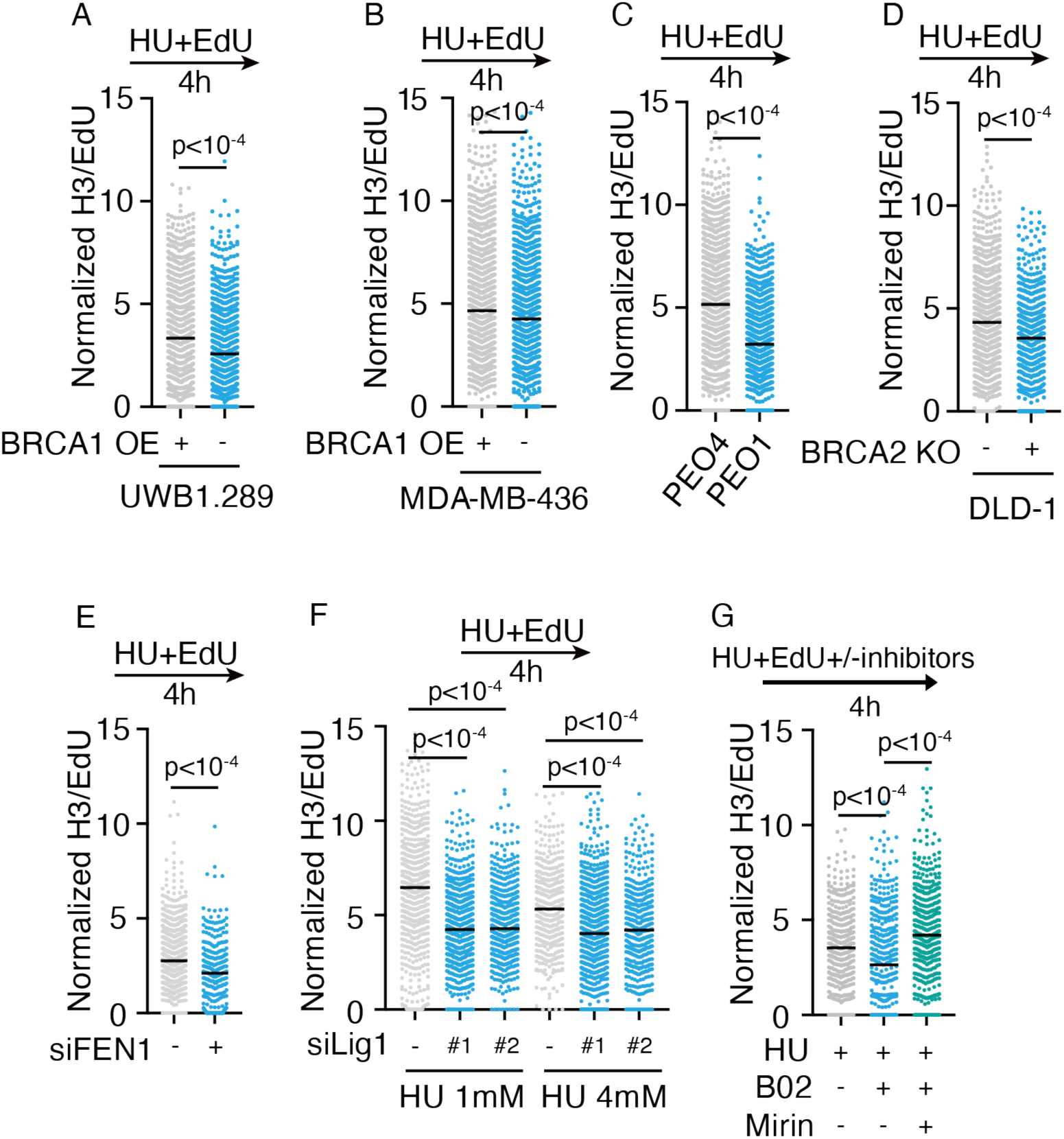
Single-stranded DNA gaps reduce nucleosome density on nascent DNA in cells. A–G) PLA for EdU and H3 in the indicated cell lines and treatments. PLA signals normalized to EdU intensity; ≥1,000 cells/sample. N ≥ 3. p-values from two-tailed unpaired t-test for A-E), Kruskal-Wallis test for F-G). UWB1.289 and MDA-MB-436 are BRCA1 deficient cells, WT BRCA1 was complemented in these cell lines. PEO1 is BRCA2 truncate mutation cell line, PEO4 is reversed back that express WT BRCA2. B02: RAD51 inhibitor, 25μM was used in this assay. Mirin: MRE11 inhibitor, 10μM used in this assay.

Defects in Okazaki fragment maturation, such as knockdown of DNA ligase 1 (LIG1) or FEN1 (Flap Endonuclease 1), combining HU treatment, also result in ssDNA gaps on the lagging strand in previous study^85^. We depleted LIG1 and FEN1 with siRNA in U2OS WT cells, and PLA signals of H3 and EdU also reduced (Figure 4E-F, Figure S5E–F). Inhibition of RAD51 triggers nucleolytic degradation of nascent DNA by nucleases such as MRE11, producing ssDNA gaps^84,86^. Therefore, we treated WT U2OS cells with RAD51 inhibitor B02 and observed low nucleosome density, and this effect was rescued by co-treatment with MRE11 nuclease inhibitor (Figure 4G, Figure S5G).

In summary, we conclude that ssDNA gaps present in post-replicative nascent DNA reduce the nucleosome density on nascent DNA.

### Single strand DNA gaps do not block nucleosome assembly in the in vitro assay

We hypothesized that single-stranded gaps in nascent DNA impair the ability to form nucleosomes. The ability of nucleosomes to form on ssDNA remains controversial. During DSB resection, the generation of long ssDNA tails/overhangs have been shown to evict nucleosomes in some studies^87–93^, whereas others have demonstrated nucleosome formation on ssDNA in vitro^94,95^. Additionally, it remains unclear whether nucleosomes can assemble on short ssDNA gaps like those found at the replication fork and what impact these short ssDNA gaps have on nucleosome stability. To determine whether nucleosomes can be assembled on DNA in the presence of ssDNA gaps, we designed a series of Widom 601 strong positioning oligonucleotides containing ssDNA gaps of various lengths (from 1–31 nts) positioned at the nucleosome dyed or the nucleosome entry/exit site (Supplemental Table 1). We then reconstituted recombinant nucleosomes containing human histones on these oligonucleotides containing ssDNA gaps using an established salt-dialysis method^96–98^ (Figure 5A). Strikingly, we observed robust nucleosome formation regardless of the ssDNA gap size or ssDNA gap position within the nucleosome (Figure 5B-D), indicating that small ssDNA gaps do not inhibit nucleosome formation *in vitro*.

**Figure 5.**
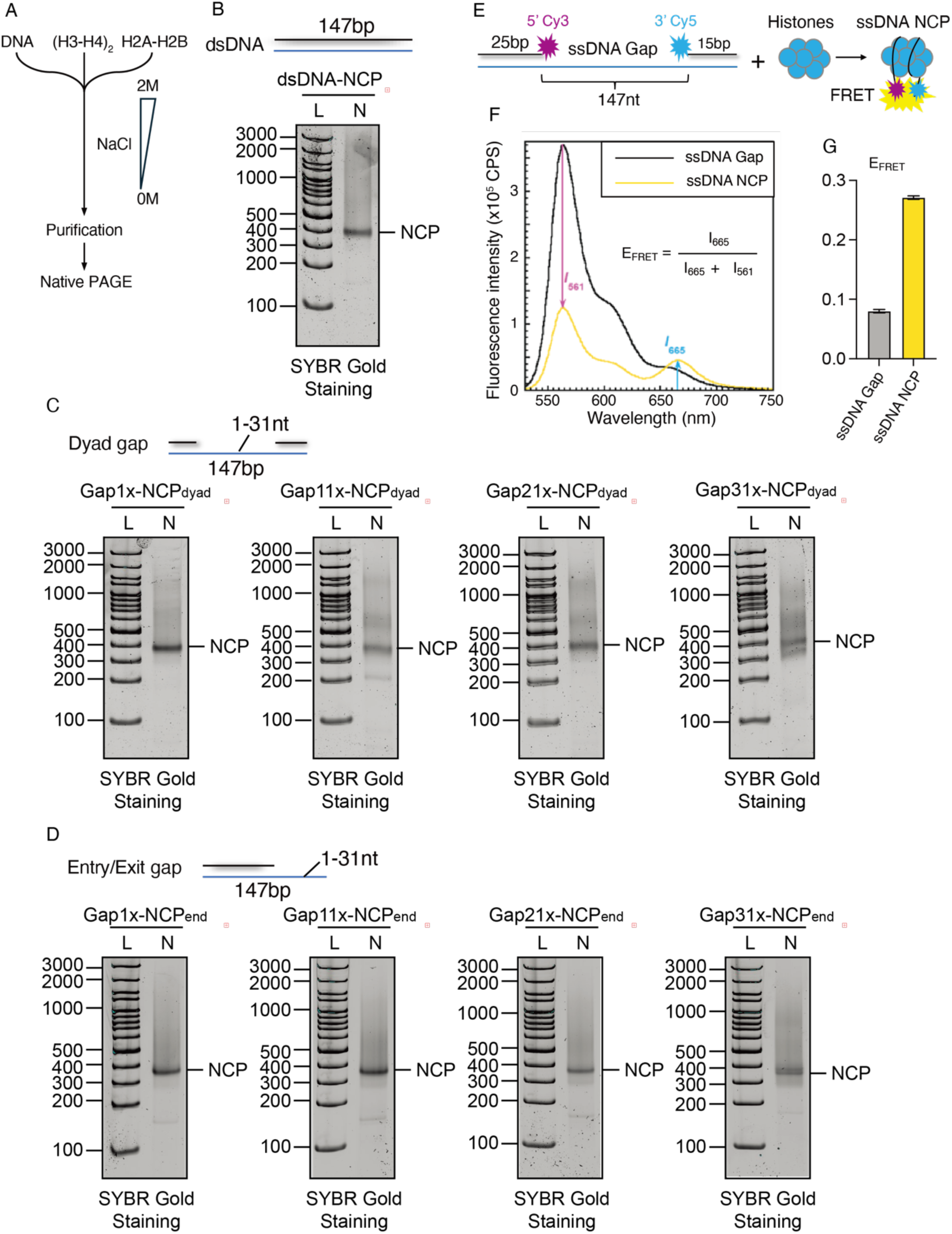
Nucleosomes can form on ssDNA gaps in vitro. A) Schematic of nucleosome formation procedure. B) Native PAGE gel image for nucleosome formation on dsDNA substrate. C) Nucleosome formation assay for dyad gap substrates. Gap size is 1-31bp as indicated. D) Nucleosome formation assay for entry/exit gap substrates. Gap size is 1-31bp as indicated. E) Schematic of the procedure to assemble a nucleosome on the 147nt ssDNA gap substrate (ssDNA Gap). Wrapping of the poly(dT)147 ssDNA around the histone octamer juxtaposes the two duplex regions and, hence, the cyanine labels, yielding a FRET signal. F) Fluorescence emission spectra obtained by excitation at 514 nm. Each spectra is the average of at least two independent spectra with the SEM shown in grey. The fluorescence emission intensities at 665 nm (Cy5 FRET acceptor fluorescence emission maximum, I_665_) and 561 nm (Cy3 FRET donor fluorescence emission maximum, I_561_) are indicated. Cy5 can be excited through FRET from Cy3 only when the two dyes are in close proximity of each other (less than ∼10 nm). This is indicated by an increase in I_665_ and a concomitant decrease in I_561_. G) Steady state FRET. The approximate FRET efficiency (*E*_FRET_) for the ssDNA Gap DNA substrate alone (ssDNA Gap) or pre-assembled in a nucleosome (ssDNA Gap Nuc) was measured at equilibrium. The height of each column (*E*_FRET_) is the average of at least 4 independent measurements with the SEM shown. FRET is only observed when a nucleosome is pre-assembled on the ssDNA Gap DNA substrate.

Having established that short ssDNA gaps (1–31 nt) at either the entry/exit or dyad regions do not markedly impair nucleosome assembly (Fig. 5B–D), we next asked whether longer gaps approaching the footprint of a full nucleosome would influence histone wrapping or stability. To address this, we generated a 147-nt ssDNA gap substrate flanked by a 25bp duplex regions with Cy3 label, and a 15bp duplex regions with Cy5 label (Figure 5E). We performed a high-resolution Cy3-Cy5 FRET (Förster Resonance Energy Transfer) assay to monitor nucleosome formation and conformational dynamics in real time (Figure 5E). When a nucleosome assembles onto a 147-nt ssDNA gap, the Cy3 and Cy5 fluorophores are brought into close proximity, resulting in a FRET signal (Figure 5F-G). Thus, the FRET signal serves as a sensitive readout for nucleosome assembly on ssDNA gaps. The steady-state FRET efficiency (E_FRET_ ≈ 0.27) confirmed the proximity of the labeled duplex arms (Figure 5F-G), consistent with a compact, fully wrapped configuration. These results demonstrate that even extensive single-stranded gaps are compatible with nucleosome formation, revealing that the histone octamer has an intrinsic ability to engage and stabilize flexible ssDNA regions. Together, these experiments (Figure 5), consistent with previous reports^95^, demonstrate that ssDNA gaps do not inherently preclude nucleosome formation in vitro. The discrepancy between the in vivo (Figure 3 and Figure 4) and in vitro (Figure 5) results suggests that the reduced nucleosome density observed in cells is likely caused by ssDNA-binding proteins, rather than by the presence of ssDNA itself (Figure S6A).

### RPA depletion does not rescue the low nucleosome density in cells

Nucleosomes reconstituted on ssDNA (ssDNA Nucs) are similar in size and morphology to those assembled on dsDNA (dsDNA Nucs) and the histone octamer core engages the wrapped DNA with similar affinities in both complexes at physiological ionic strength^94,95,99–102^. However, ssDNA Nucleosomes are more disordered and dynamic^95,100^, suggesting that regions of the ssDNA wrapped around a histone octamer core are dynamically exposed. The major ssDNA-binding protein complex, Replication Protein A (RPA), is highly abundant (10 μM range) in the nucleoplasm ^103–105^ and has the highest ssDNA affinity (*K*_D_ ∼ pM to nM range) of any ssDNA-binding protein/protein complex that has been identified in human cells. Each RPA engages ∼20-33 nt of ssDNA, extending the bound sequence into a linear conformation and increasing its bending 2 to 3-fold ^106–117^.

To determine whether RPA can substitute for nucleosomes on ssDNA, we performed parallel in vitro and in vivo experiments. In vitro, we compared the FRET signal of RPA binding to the ssDNA gaps only, and confirmed that RPA binding on ssDNA gaps can’t cause FRET signal (ssDNA gaps with RPA E_FRET_=0.075±0.001, ssDNA gaps only E_FRET_=0.08±0.001, Figure S6B). Next, we incubate pre-assembled ssDNA nucleosomes with physiological concentrations of RPA, and we observed a time-dependent decline in FRET efficiency (E_FRET_ decreasing from 0.25±0.001 to 0.119±0.001; Figure S6C).

Furthermore, E_FRET_ values decrease upon the addition of RPA down to an equilibrium value (E_FRET_ = 0.119 + 0.001, Figure S6C) that is noticeably higher than the equilibrium E_FRET_ values observed for the ssDNA Gap DNA substrate in the absence (0.080 + 0.001) and presence of RPA (0.075 + 0.001) (Figure S6B). It cannot be discerned whether the elevated equilibrium value (Compare the E_FRET_ in Figure S6C to E_FRET_ in Figure S6B) is due to indirect effects of the fully unwrapped histone octamer on the RPA-ssDNA filament or the presence of a minor population of stable, fully wrapped ssDNA nucleosomes or partially wrapped ssDNA Nucleosomes. Nonetheless, the clear decrease in E_FRET_ observed upon the addition of RPA suggests that ssDNA readily unwraps from around histone octamer cores and the exposed sequences are engaged and stabilized by RPA.

We performed immunofluorescence staining for RPA in FRKO cells to corroborate the in vitro findings, since we observed increased levels of RPA2 and its phosphorylation, consistent with elevated ssDNA accumulation in FRKO cells upon HU treatment (Figure S5D and S6E). To test whether elevated RPA contributes to the reduced nucleosome density, we conducted a PLA assay for H3 and EdU in RPA1/2-depleted U2OS cells (Figure S6F). However, depletion of RPA1 and/or RPA2 did not rescue the low nucleosome density in fork-reversal-deficient cells. These results suggest that RPA does not directly evict nucleosomes from ssDNA in cells, or that other ssDNA-binding factors may perform this function when RPA is depleted.

### Transcription reduces nucleosome density in cells

Since we observed reduced chromatin compaction marks, such as H3K9me2/3 (Figure 2) and lower nucleosome density in FRKO cells (Figure 1), we hypothesized that chromatin decompaction may increase DNA accessibility to enzymes such as RNA polymerase II (Pol II). Conversely, transcription elongation and R-loop formation could further disrupt nucleosomes on nascent DNA. Thus, we proposed that RNA Pol II–mediated transcription on nascent DNA and low nucleosome density may reinforce each other.

To test this hypothesis, we first examined RNA Pol II occupancy on nascent DNA using a PLA assay targeting EdU and the major RNA Pol II subunit RPB1 (Figure S7A). We observed a significant increase in PLA signal in FRKO cells, indicating elevated Pol II association with nascent DNA. As a control, treatment with Triptolide (TPL), a potent RNA Pol II inhibitor that induces proteasome-dependent degradation of Pol II, dramatically reduced the RPB1 PLA signal, confirming the signal specificity for RNA Pol II (Figure S7A).

To determine whether increased transcription contributes to the reduced nucleosome density, we performed a PLA for H3 and EdU in TPL-treated WT and FRKO cells. TPL treatment significantly increased nucleosome density in both WT and FRKO cells, suggesting that Pol II activity on nascent DNA can reduce nucleosome density (Figure S7B). However, TPL treatment did not fully restore nucleosome density in FRKO cells to the levels in WT U2OS cells, indicating that transcription alone cannot account for the nucleosome loss in FRKO cells, and additional fork reversal dependent mechanisms likely contribute to this phenotype.

### PARylation suppresses nucleosome assembly on ssDNA gaps in fork reversal deficient cells

PARylation is known to alter chromatin structure and orchestrate DNA repair^118–124^. We observed the increase in PARylation in FRKO cells upon HU treatment (Figure 6A), suggesting increased ssDNA gaps in FRKO cells. To determine if PARP1-mediated PARylation impacts nucleosome density, we depleted PARP1 in FRKO cells via siRNA. PARP1 depletion reduced total level of PARylation and partially rescued nucleosome density despite persistent ssDNA gaps (Figure 6B-D). Inhibition of PARG, the enzyme that hydrolyzes PAR chains, increased PARylation and decreased nucleosome density in WT cells, further supporting the conclusion that PARylation reduces nucleosome density in cells (Figure 6E and Figure 6F). Interestingly, treatment with the PARP1/2 inhibitor Olaparib reduced PARylation in WT cells after HU treatment (Figure 6G), but nucleosome density still decreased (Figure 6H). This indicates that PARP inhibition alone can lower nucleosome density on nascent DNA. In FRKO cells, Olaparib similarly reduced PARylation but nucleosome density remains low, reinforcing that PARP1 inhibition itself causes this effect. Together, these findings indicate that ssDNA activates PARP1-mediated PARylation, which in turn reduces nucleosome density on nascent strand DNA. PARP1 trapping by Olaparib can also reduce nucleosome density on nascent DNA.

**Figure 6.**
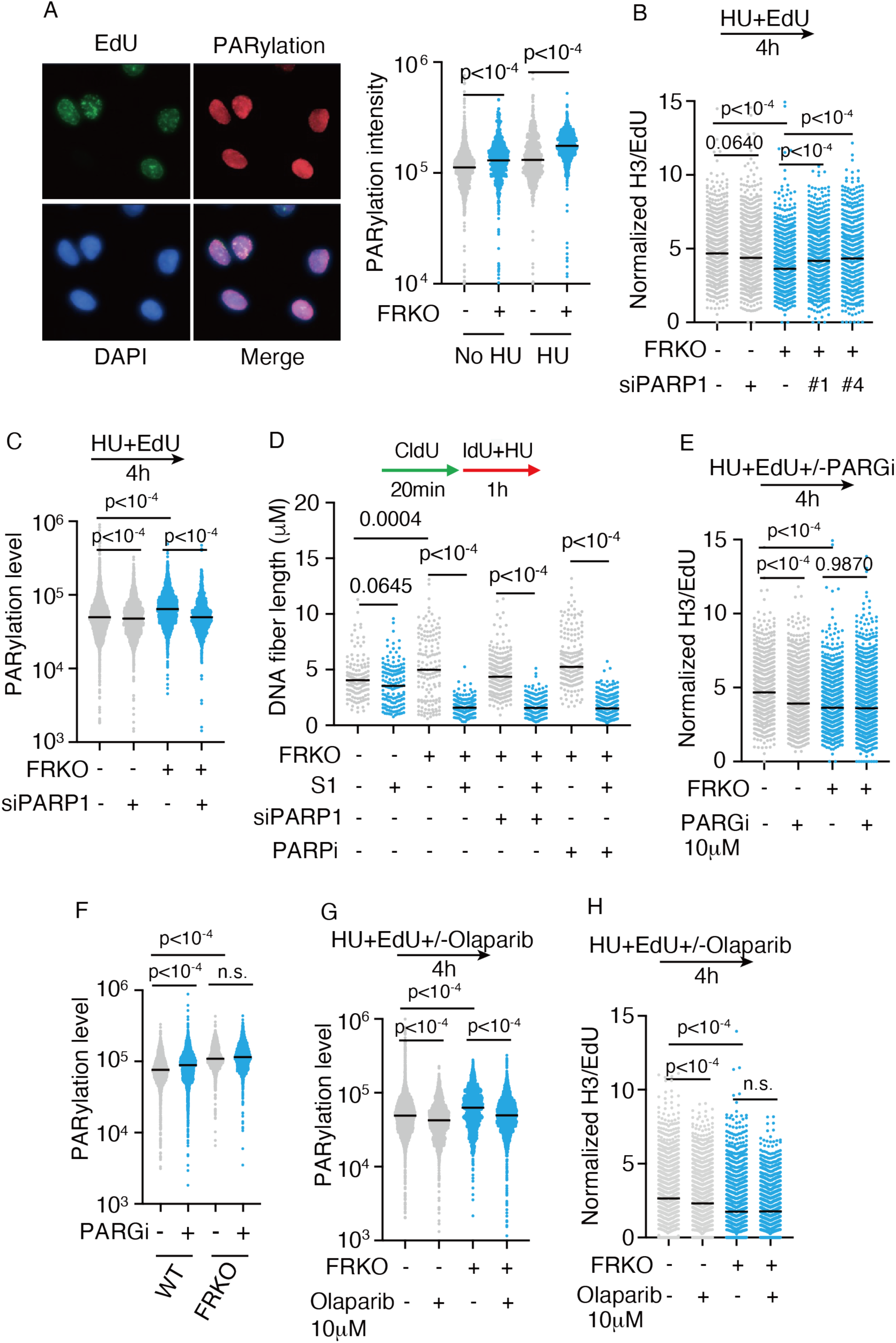
PARylation reduces nucleosome density in fork reversal-deficient cells. A) Immunofluorescence of PARylation in WT and FRKO cells. Cells were labeled with EdU for 10 min in no HU samples, or EdU with HU for 4 h. Integrated PARylation intensity quantified. N = 3. p-values from Kruskal-Wallis test. B) PLA for EdU and H3 under indicated conditions. PLA normalized to EdU intensity; ≥1,000 cells/sample. N = 9. p-values from Kruskal-Wallis test. C) Immunofluorescence of PARylation in indicated conditions. N = 3. D) S1 nuclease DNA fiber assay to detect ssDNA gaps. N = 3. p-values from Kruskal-Wallis test. E) PLA for EdU and H3 in indicated conditions, quantified as in (B). N ≥ 7. F-G) Immunofluorescence for PARylation in the indicated conditions. N = 3. H) PLA for EdU and H3 in indicated conditions, quantified as in (B). N ≥ 7.

### HMCES–DPC inhibits nucleosome assembly on ssDNA

We sought additional factors that might bind ssDNA and inhibit nucleosome assembly. From our iPOND-MS data, we noticed that in fork reversal–deficient cells, APOBEC3B levels at replication forks increased by ∼1.5-fold (Figure 7A), chromatin-bound APOBEC3B increased by 3–5-fold (Figure S6A). APOBEC3B can deaminate cytosine residues, which are subsequently processed by glycosylases to generate AP sites^125^. HMCES, a PCNA-binding protein, has been shown to form covalent DNA–protein crosslinks (DPCs) at AP sites on ssDNA^126–132^. We observed a modest increase in levels of HMCES in iPOND-MS (Figure 7A), and a significant increase in HMCES-DNA crosslink by RADAR assay (Figure 7B).

**Figure 7.**
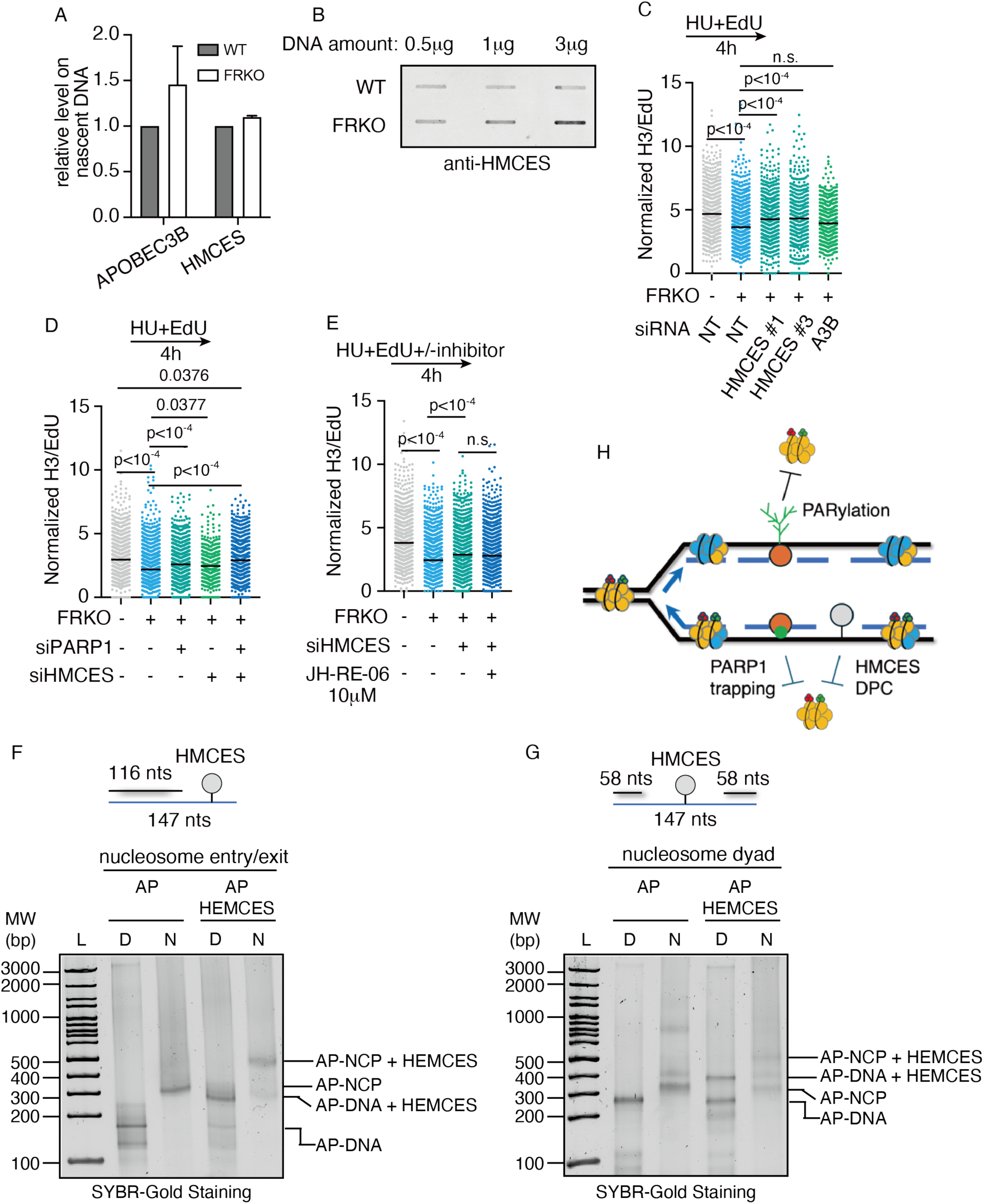
HMCES-DPC at ssDNA gaps reduces nucleosome density in fork reversal-deficient cells. A) APOBEC3B and HMCES levels on nascent DNA measured by iPOND-SILAC-MS. N = 4. B) RADAR assay detecting HMCES-DPC in WT and FRKO cells. N = 3. C–E) PLA for EdU and H3 in indicated conditions (as in Figure 1B). N ≥ 7. p-values from Kruskal–Wallis test. F–G) Nucleosome reconstitution assays showing nucleosomes fail to form on dyad ssDNA gaps but can form at side gaps. Gels show NCP bands. AP-DNA represents abasic-site DNA; AP-NCP represents nucleosomes assembled on abasic DNA. L represents ladder, D is DNA, N is nucleosome. H) Model schematic of how ssDNA adducts interfere with nucleosome formation at gaps.

We hypothesized that in FRKO cells, the high abundance of ssDNA on nascent DNA permits excess APOBEC3B activity, leading to the generation of AP sites that subsequently form HMCES–DPCs. These DPCs could represent a steric hindrance that blocks nucleosome assembly. To test this hypothesis, we depleted HMCES or APOBEC3B using siRNA and measured nucleosome density on nascent DNA by PLA (Figure 7C, Figure S8B–C). HMCES depletion restored nucleosome density in fork reversal–deficient cells, whereas APOBEC3B depletion only mild rescue the low nucleosome density, but not statistically significant (Figure 7C), likely because AP sites can also form spontaneously on ssDNA independent of APOBEC3B. Co-deletion of PARP1 and HMCES additively rescued nucleosome density in fork reversal–deficient cells (Figure 7D). HMCES loss is also known to activate TLS (translesion synthesis) for post-replication repair^127^, which in turn repair ssDNA gaps^67,80^. We next asked whether the rescue of nucleosome density upon HMCES depletion was mediated by TLS-dependent gap repair. Inhibition of TLS using the REV1 inhibitor JH-RE-06 in HMCES-deficient cells did not reduce nucleosome density (Figure 7E), demonstrating that HMCES–DPCs themselves, rather than TLS activation, block nucleosome assembly.

### HMCES-DPC at dyad position inhibits nucleosome formation *in vitro*

To determine if a HMCES-DPC can block nucleosome assembly *in vitro*, we designed two Widom 601 strong positioning oligonucleotides containing a 31 nt ssDNA gap positioned at the nucleosome dyed or the nucleosome entry/exit site, with an uracil located in the ssDNA region (Figure S8D-E). These DNA substrates were treated with uracil DNA glycosylase to generate an AP-site, and the HMCES-AP-site DPC stabilized using sodium borohydride (Figure S8F-G)^127,128,133^. We then attempted to reconstitute recombinant human nucleosomes on these oligonucleotides containing the HEMCES-DPC using a salt-dialysis method (Figure 5A)^96–98^. Notably, we observed robust nucleosome formation when the HMCES-DPC was positioned at the nucleosome entry/exit site (Figure 7F). In contrast, we observed minimal nucleosome formation when the HMCES-DPC was positioned in the DNA near the nucleosome dyad, indicating that HMCES represents a steric hindrance to nucleosome formation (Figure 7G). We hypothesize that elevated nucleosomal DNA dynamics at the entry/exit site compared to the dyad enables the nucleosome to accommodate the HMCES-DPC at this location^134^. Together, this data shows that a HMCES-DPC can directly impair nucleosome formation, though the ability to accommodate a HMCES-DPC will likely be dictated by the location of the HMCES-DPC in relation to the histone octamer during histone deposition on ssDNA gaps.

Together, these findings demonstrate that in FRKO cells, PrimPol-mediated repriming generates abundant ssDNA gaps, which activate PARP1-dependent PARylation and promote HMCES–DPC formation. At least three mechanisms: Transcription, PARylation and HMCES–DPC formation act in concert to inhibit nucleosome assembly on ssDNA gaps, thereby reducing parental histone recycling and lowering nucleosome density on nascent DNA (Figure 7H).

## Discussion

### Fork reversal is essential for both genome stability and epigenome stability

In this study, we demonstrate that fork reversal is critical not only for genome stability, but also for epigenome stability during DNA replication under stress. We propose a mechanistic model in which fork reversal suppresses ssDNA gap formation, thereby promoting parental histone recycling and nucleosome assembly on nascent DNA to preserve chromatin states. In the absence of fork reversal, PrimPol-mediated repriming generates abundant ssDNA gaps, which trigger PARP1-dependent PARylation and HMCES–DPC formation (Figure 6 and Figure 7). These events hinder nucleosome deposition and lead to the loss of parental histone modifications (Figure 2 and Figure 3). Together, our work establishes a mechanistic paradigm linking replication fork intermediates (fork reversal and ssDNA) to faithful epigenetic inheritance, reframing replication stress from a purely DNA-centric problem to a driver of epigenetic instability with broad implications for human disease.

### Loss of parental-specific histone marks on nascent DNA in FRKO cells reflects impaired parental histone recycling

A previous elegant study showed that HU treatment promotes accumulation of H3K9me2/3 at stalled forks, thereby helping protect stalled replication structures{Gaggioli, 2023 #157}. An important question is whether, in fork-reversal–deficient (FRKO) cells, the reduced abundance of parental-specific histone modifications on nascent DNA (e.g., H3K9me2/3) primarily reflects loss of parental histones, a failure to induce/enrich these marks at stalled forks, or a combination of both. Our data support the conclusion that the decrease at least partially reflects impaired parental histone recycling for two reasons. First, HU-induced enrichment of H3K9me2/3 at stalled forks is ATR-dependent and has been reported to occur independently of fork reversal{Gaggioli, 2023 #157}. Consistent with this, we observed even stronger ATR activation in FRKO cells (Figure S6E), suggesting that an inability to activate ATR is unlikely to explain the reduced parental-mark signal in our system. Second, we detected a broad reduction across multiple parental histone–associated signatures, not limited to H3K9me2/3, including H4K20me2, H3K79me2, and the enrichment of histone variants such as CENP-A and macroH2A.

In summary, these findings indicate that fork reversal deficiency leads to a loss of parental histone information on nascent DNA, consistent with defective parental histone recycling. However, we cannot fully exclude the possibility that FRKO cells also exhibit a reduced capacity to establish or enrich specific marks under HU compared with WT cells. These mechanisms are not mutually exclusive, and both impaired parental histone recycling and altered mark induction may contribute to the phenotype observed under sustained HU treatment.

### How parental histone loss in FRKO cells

Our results suggest that PrimPol repriming–mediated ssDNA gaps contribute to reduced nucleosome density on nascent DNA. However, our data do not fully exclude an alternative possibility: that FRKO cells experience impaired histone delivery during DNA synthesis due to altered usage of histone-chaperone–coupled replicative polymerases. Current evidence does not support a direct histone chaperone activity for PrimPol. Thus, one speculative model is that, in FRKO cells, PrimPol extends nascent DNA in contexts where replicative polymerases with tighter histone-coupled synthesis (e.g., Pol ε) would normally operate, resulting in DNA extension that is relatively uncoupled from histone deposition. Although we did not detect an obvious change in Pol ε abundance by iPOND-MS (Figure 3A), we can’t fully exclude this possibility.

Based on our findings together with prior studies, we conclude that ssDNA is not intrinsically incompatible with nucleosome assembly: from a biophysical standpoint, ssDNA can support the formation of ssNUCs. The critical unresolved issue is the temporal and spatial order of ssDNA formation relative to ssDNA-associated adducts, particularly DPCs and PARylation. Because we cannot determine when and where DPCs and PARylation arise with respect to ssDNA gaps, our data are consistent with two alternative, non–mutually exclusive models: 1) Pre-existing ssDNA regions are present in the genome prior to fork passage and are already decorated with DPCs and/or PARylation. In this scenario, as replication proceeds through these regions, parental histones would be unable to efficiently recycle onto the ssDNA substrate, resulting in an immediate deficit in nucleosome density on nascent DNA. 2) Replication itself generates abundant ssDNA gaps; ssNUCs can initially form on or near these ssDNA regions, but subsequent accumulation of DPCs and/or PARylation destabilizes these structures and promotes their eviction, thereby reducing nucleosome density.

Although our experiments show that HU treatment is associated with decreased nucleosome density, they do not definitively exclude the first model. Notably, HU exposure in our assays is sustained for ∼4 hours, which could allow the establishment of pre-existing ssDNA lesions and associated adducts before or during replication, making it difficult to unambiguously assign causality and order of events.

Previous work has elegantly shown that under replication stress, ASF1 can sequester parental histones during fork stalling and facilitate their recycling upon fork restart^136^. Here, we focused on parental histone recycling during ongoing replication under stress and observed a pronounced loss of parental histones in FRKO cells. One possibility is that excessive ssDNA gaps in FRKO cells increase the frequency of parental histone disassembly, which in turn may overwhelm the buffering capacity of ASF1, ultimately resulting in defective parental histone recycling and reduced nucleosome density on nascent DNA.

### Parental histones are lost during replication under stress in FRKO cells

These parental marks and variants (H4K20me2, H3K9me3, MacroH2A.1/2, CENP-A) are generally repressive features that promote chromatin compaction and transcriptional silencing. H3K9me3-marked heterochromatin, in particular, is enriched at repetitive genomic regions, where it suppresses transcription^137^. Thus, the loss of parental histones in FRKO cells likely results in the derepression of repetitive elements. Prior studies in mouse embryonic stem cells have shown that disrupting parental-histone recycling via depletion of histone chaperones activates transposon elements, thereby triggering inflammatory and immune signaling and impairing differentiation and cell viability^49,50^. SMARCAL1 loss has been implicated increasing the response to immunotherapy through activating cGAS-STING pathway, and suppressing PD-L1 express^138^. Our study provided extra rationale that fork reversal can be targeted for immunotherapy since its loss may activate repeat genes, therefore producing antigens for activating cytokines.

### ssDNA gaps reduce the nucleosome density on nascent DNA

Notably, all our cellular assays are EdU labeling-based, as we use EdU label nascent DNA, such as PLA, iPOND, and ChOR-seq. Therefore, these assays may not resolve ssDNA-nucleosomes beyond the resolution distance of these assays. This assumption may explain the inconsistency between in vivo and in vitro assays. However, the spatial resolution of PLA is approximately 40 nm (∼100 bp/nt), while the DNA fragments analyzed by iPOND and ChOR-seq are typically 100–400 bp/nt in length^72,139^. Previously published electron microscopy images showed that most ssDNA gap sizes on nascent DNA are less than 300bp^53,66,67,80,140^, in very rare cases can reach up to 1000 nt. The ssDNA-nucleosomes are even closer to the EdU in dsDNA spatially after nucleosome assembly (Figure 5E). Therefore, we hypothesize that cellular assays such as PLA, ChOR-seq, and iPOND should have enough resolution to detect most of the ssDNA-nucleosome occupancy, as long as it exists. However, developing higher-resolution assays will be necessary to further confirm whether nucleosomes bind ssDNA directly in cells in the future.

### ssDNA gap events that inhibit histone assembly

We conclude that PARylation, transcription, and DPC are three events that reduce nucleosome density. PARP1-mediated PARylation promotes chromatin relaxation via multiple mechanisms: 1) the negatively charged poly(ADP-ribose) (PAR) chains neutralize histone positive charges and weaken histone–DNA interactions^122^; and 2) PAR chains act as docking platforms to recruit DNA repair enzymes and chromatin remodelers, such as ALC1 (CHD1L)^119,120,141–143^, which remodels nucleosomes. Together, these mechanisms enhance chromatin accessibility, exposing ssDNA regions to repair factors. Surprisedly, we found that PARP inhibitor Olaparib treatment reduce the nucleosome density on nascent DNA, which is distinct from PARP1 depletion. We hypothesize that, PARP1 depletion removes PARylation and leaves ssDNA gaps relatively “clean,” so the nucleosome can form on the ssDNA gaps. However, PARP inhibitor blocks PARylation and trap PARP1 on DNA, since PARylation normally promotes PARP1 release^145,146^. We therefore propose that trapped PARP1 directly displaces nucleosomes. This idea is consistent with recent findings showing that PARP1 inhibition leads to histone eviction from replication fork ^147^.

We examined the role of HMCES-DPCs, which were significantly increased in FRKO cells (Figure 7B). HMCES depletion rescued the low nucleosome density in FRKO cells, as shown by PLA (Figure 7C), and this was further confirmed by in vitro reconstitution assays (Figure 7F-G). Surprisingly, HMCES-DPCs impaired nucleosome assembly only when positioned near the nucleosome dyad (Figure 7G), not at the entry or exit sites (Figure 7H). However, we cannot rule out inhibitory effects of entry/exit DPCs in cells. This is because the discrepancy between in vivo and in vitro system: (1) our reconstitution assays involve single nucleosomes, where ssDNA gaps at the entry/exit sites are flexible and may minimize DPC effects; (2) the Widom 601 sequence was specifically selected for nucleosome formation assay in previous studies, which has high histone-binding affinity, which could overcome HMCES-DPC–mediated inhibition; and (3) in cells, HMCES-DPCs can undergo additional processing, such as ubiquitination or proteolytic degradation, triggering downstream repair cascades that may collectively reduce nucleosome density^127,144^. These complex in vivo events are not captured by simplified in vitro systems.

We cannot exclude other ssDNA binding proteins, or any ssDNA-related events that also compromise nucleosome formation. Therefore, we conclude that PARylation, RNA Pol II, and DPC are the least events that inhibit histone assembly.

## Materials and Methods

### Cell lines

U2OS and MDA-MB-436 cells were cultured in DMEM with 9% fetal bovine serum (FBS) (Bio-techne). PEO4, PEO1, and DLD-1 cells were cultured in RPMI-1640 with 9% FBS. UWB1.289 cells were cultured in a 1:1 mixture of RPMI-1640 medium and Mammary Epithelial Growth Medium (MEGM), supplemented with FBS and antibiotics. For iPOND-MS, U2OS cells were cultured in SILAC-compatible DMEM (Thermo Fisher Scientific A33822) with 10% dialyzed FBS (R&D Systems S12850H) and either isotopically light or heavy 13C, 15N lysine, and 13C,15N arginine (Cambridge Isotope Laboratories)^70^. Cells were cultured at 37°C and 5% CO2 with humidity. All cell lines were regularly tested for mycoplasma and verified using short tandem repeat profiling.

### Genome editing

PrimPol knockout was generated using CRISPR-Cas9 with the SpCas9(BB)-2A-Puro (PX459) V2.0 plasmid (Addgene #62988) as described^148^. The single guide RNA (sgRNA) sequence was 5’-CTGCCAAGCAAGTCAAGAGC-3’. Transfected cells were selected with puromycin, and single-cell clones were isolated and validated by sequencing and immunoblotting.

### SILAC and Label free iPOND-mass spectrometry

iPOND-SILAC MS was performed as described previously^70^. Briefly, EdU-labeled and indicated drug-treated cells were crosslinked with 1% formaldehyde, quenched with glycine, permeabilized in 0.25% Triton X-100. Heavy-labeled and light-labeled cells were combined 1:1 prior to the click reaction. DNA-protein complexes were captured with Dynabeads MyOne Streptavidin C1 (Thermo Fisher 65002), washed, and eluted with elution buffer (25mM biotin, 0.4% SDS, 50mM Tris-HCl (pH 8.5)) at 65°C. Supernatant, containing eluted fractions, were incubated at 95°C to reverse crosslinking. For histone modification analysis, lysine were derivatized with 2.5% propionic anhydride (Sigma-Aldrich 8.00608) in 50 mM ammonium bicarbonate (pH 7.5). Subsequently, proteins were digested with 200 ng trypsin (Thermo Fisher 90057) in 50 mM ammonium bicarbonate (pH7.5) overnight. Peptides were desalted using C18 Stagetips (reversed-phase resin) and carbon Top-Tips (Glygen TT1CAR), speed-vacuumed dried, and stored at −20°C until MS analysis.

### PrimPol complementation

PrimPol WT, AxA, and CH mutants were subcloned from pcDNA3.1/nV5-DEST to pLNCX2 using forward primer: 5’-CAGATCTCGAGATGAATAGAAAATGGGAAGCAA-3’ and reverse primer: 5’-CTGCGGCCGCCTCTTGTAATACTTCTATAATTAG-3’. pLNCX2-Primpol constructs and pVSV-G were co-transfected into GP2-293 cells using Polyethylenimine (PEI). Twenty-four hours post-transfection, the culture medium was replaced with fresh medium, and viral supernatants were harvested at 48 and 72 hours. The pooled viral supernatants were subsequently used to infect U2OS Primpol knockout #206 cells. Stable cell lines were established by selection with 100 µg/ml G418.

### DNA combing assays

Cells were labeled with the nucleoside analogs as indicated in the figures and figure legends. For ssDNA gap detection, S1 nuclease digestion assay was performed as previously described^66^. DNA combing was performed using a DNA combing instrument as previously described^54^.

### Immunofluorescence

For analysis of proteins in the insoluble/chromatin fraction, cells were pre-extracted with 0.5% Triton X-100 and subsequently fixed in 4% paraformaldehyde. Slides were blocked with 1% goat serum in PBST (0.1% triton-X100) and incubated with primary antibodies. EdU incorporation was visualized by click chemistry with an Alexa Fluor 488-conjugated azide. Immunofluorescent images were obtained and analyzed by ImageXpress microscopy and software (Molecular Devices).

### PLA assay

Proximity ligation assays (PLAs) were performed using the Duolink kit (Sigma DUO92002, DUO92004, DUO92008) according to the manufacturer’s instructions, with the following modifications as described previously ^71^. Cells were pre-extracted and fixed as described in the immunofluorescence method, then blocked with 3% BSA and processed by click chemistry with biotin-azide and Alexa Fluor 488-conjugated azide (9:1 ratio). The following primary antibody pairs were used: rabbit H2A (CST 12349) (1:100 dilution), H3 (Abcam ab1791) (1:200 dilution), H3K9ac (Millipore 07-352) (1:100 dilution), H3K9me3 (CST 13969) (1:100 dilution), H4 (CST 13919) (1:100 dilution); mouse biotin (Millipore B7653) (1:200 dilution) and mouse H2B (Abcam ab52484) (1:100 dilution), H3K9me2 (Abcam ab1220) (1:100dilution) and rabbit biotin (CST 5597) (1:100 dilution). Images were obtained by ImageXpress Pico and analyzed by MetaXpress (Molecular Devices).

### CHOR-Seq

ChoR-seq was performed as previous described^72^. Briefly, WT and FRKO U2OS cells (n = 3 biological replicates each) were labeled with 10 µM EdU for 15 min for no HU sample. For HU treated sample, the cells are treated with 10uM EdU and 1 mM HU (Sigma H8627) for 4 h before fixation with the truChIP kit (Covaris 520127). Chromatin was sonicated to 100–500 bp, and EdU-labeled Drosophila S2 chromatin was added at 0.05% (w/w) as a spike-in for normalization. H3 ChIP was performed with anti-H3 (10 µg, Abcam ab1791). Immunoprecipitated DNA was processed with the NEBNext Ultra II DNA Library Prep Kit (NEB E7645S), indexed, and sequenced on an Illumina NovaSeq platform.

### Data processing

All sequencing reads were mapped to the H. sapiens reference genome (hg38) and D. melanogaster reference genome (BDGP6) using Bowtie2 (v2.5.4) with the default parameters^149^. Only uniquely mapping reads were kept, and PCR duplicates were removed using Markup from Samtools (v1.17) (https://www.htslib.org/doc/samtools-markdup.html). Genome occupancy was calculated using bamCoverage from Deeptools (v3.5.5)^150^ with standard parameters in 100 bp intervals, normalized to relative reads per million (RRPM) using D. melanogaster reads as described previously^151^, and visualized by the Integrative Genomics Viewer (IGV) (v2.18)^152^. To generate a common H3 peak dataset, peak calling in each sample were preformed using MACS2 (v2.2.7)^153^ with parameters of “--broad-cutoff 0.1” and merged into a consensus region set using the Intersect function in Bedtools (v2.26.0)^154^. Subsequently, for H3 overlapped peaks in RRPM-normalized datasets were exported with the computeMatrix tool from Deeptools, and the enrichment patterns were visualized using the plotProfile and plotHeatmap function in DeepTools. Lastly, using the Intersect function in bedtools to get the H3 distribution on genome repeat elements and visualize in R. All datasets were deposited in the NCBI GEO: GSE000000.

### Preparation of oligonucleotides for recombinant nucleosomes

All oligonucleotides were synthesized by Integrated DNA Technologies (Coralville, IA). A complete list of oligonucleotides used to make recombinant nucleosomes can be found in Supplementary Table 1. Each oligonucleotide was resuspended to a final concentration of 100 µM in a buffer containing 10 mM Tris (pH-7.5) and 1 mM EDTA. Complimentary oligonucleotides (see Supplementary Table 1) were mixed to a final concentration of 10 µM and annealed by heating to 95°C for 2 minutes and cooling to 10 °C at a rate of -5 °C/min. The annealed oligonucleotides were stored short-term at 4 °C prior to nucleosome reconstitution.

### Radar Assay

Cells were lysed in Radar buffer (4M Guanidine thiocyanate, 1% Sarkosyl, 2% Triton X-100, 1% 1,4-dithioerythritol, 100 mM Sodium Acetate pH5.0, 20 mM Tris pH 8.0, 20 mM EDTA pH8.0; adjusted to pH 6.5). Genomic DNA was sonicated and precipitated with ethanol, resuspended in 8 mM NaOH, and quantified using Qbit fluorometry. Samples were then subjected to slot-blot onto nitrocellulose membrane, which were subsequently immunoblotted with HMCES primary antibody (Cell Signaling #17636) and corresponding secondary antibody.)

### RNA interference

All siRNA transfections were performed using DharmaFECT reagents (Horizon Discovery T-2001-03) according to the manufacturer’s instructions. Experiments were performed 2 days after transfection. Qiagen AllStars Negative Control Nontargeting (NT) siRNA was used as a non-targeting control. Gene-targeting siRNA sequences are provided in the supplementary table

### Purification of recombinant human histones

The genes encoding human histone H2A (UniProt identifier: P0C0S8), H2B (UniProt identifier: P62807), H3 C110A (UniProt identifier Q71DI3), and H4 (Uniprot identifier: P62805) were cloned into a pET3a expression vector. Histone H2A, H3, and H4 were individually transformed and expressed in BL21 (DE3) pLysS competent *E. coli* cells (New England BioLabs). Histone H2B was transformed and expressed in BL21-CodonPlus (DE3)-RIPL *E. coli* cells (Agilent). The histones were grown in M9 minimal media (33.5 mM Na_2_HPO_4_•7H2O, 220 mM KH_2_PO_4_, 100 mM NaCl, 200mM NH_4_Cl, pH 7.2) supplemented with 0.4% glucose (w/v), 2 mM MgSO_4_, 0.2 mM CaCl_2_ and a 1% vitamin cocktail. The histones were grown at 37 °C to an OD600 of 0.4, and histone expression induced with a 0.4 mM (histone H2A, H2B, and H3) or 0.3 mM (histone H4) isopropyl-β-D-thiogalactoside (IPTG) for 3 hours (histone H3 and H4) or 4 hours (histone H2A and H2B). The cells were harvested via centrifugation and cell pellets stored long term at −80 °C. The individual histones were purified from inclusion bodies using well-established methods^96–98^. In brief, the histone cell pellets were lysed via sonication in a buffer containing 50 mM Tris-HCl (pH 7.5), 100 mM NaCl, 1 mM benzamidine, 1 mM DTT, and 1 mM EDTA and the cell lysate clarified via centrifugation. The resulting pellet was washed three times with a buffer containing 50 mM Tris-HCl (pH 7.5), 100 mM NaCl, 1 mM benzamidine, 1 mM DTT, 1 mM EDTA, and 1% Triton X-100 and washed a final time with a buffer containing 50 mM Tris-HCl (pH 7.5), 100 mM NaCl, 1 mM benzamidine, 1 mM DTT, and 1 mM EDTA. The histones were extracted from inclusion bodies under denaturing condition (6 M Guanidinium-HCl), dialyzed into 8 M Urea, and purified using anion-exchange and cation-exchange chromatography under gravity flow. The purified histones were dialyzed five times against H_2_O, lyophilized, and stored long-term at −20 °C.

### Preparation of H2A/H2B Dimer and H3/H4 Tetramer

Prior to nucleosome assembly, H2A/H2B dimers and H3/H4 tetramers were prepared using established methods ^96–98^. Each individual lyophilized histone was resuspended in a buffer containing 20 mM Tris-HCl (pH 7.5), 6 M Guanidinium-HCl, and 10 mM DTT. To refold the H2A/H2B dimer, H2A and H2B were mixed in a 1:1 molar ratio and dialyzed three times against a buffer containing 20 mM Tris-HCl (pH 7.5), 2 M NaCl, and 1 mM EDTA at 4 °C. To refold the H3/H4 tetramer, H3 and H4 were mixed in a 1:1 molar ratio and dialyzed three times against a buffer containing 20 mM Tris-HCl (pH 7.5), 2 M NaCl, and 1 mM EDTA at 4 °C. The refolded H2A/H2B dimer and H3/H4 tetramer were subsequently purified using a HiPrep Sephacryl S-200 16/60 HR gel filtration column (Cytiva) in a buffer containing 20 mM Tris-HCl (pH 7.5), 2 M NaCl, and 1 mM EDTA. Pure fractions containing H2A/H2B dimers and H3/H4 tetramer were combined and stored long term at −20 °C in a 50% glycerol slurry.

### Nucleosome assembly and purification

All nucleosomes were prepared using a well-established salt-dialysis method ^96–98^, with minor modifications. In brief, DNA, H2A/H2B dimer, and H3/H4 tetramer were mixed at a ratio of 1:2.2:1:1, respectively, in a buffer containing 20 mM Tris-HCl (pH 7.5), 2 M NaCl, 1 mM EDTA, and 1mM DTT. Nucleosomes were assembled via a stepwise decrease in NaCl concentration over a 24-hour period - from 2 M NaCl to 1.5 M NaCl, to 1.0 M NaCl, to 0.5 M NaCl, to 0.25 M NaCl, and to 0 M NaCl. The assembled nucleosomes were then purified via ultracentrifugation for 42 to 44 hours at 4 °C using a 10% - 40% sucrose gradient. Final nucleosome purity and homogeneity was assessed by native PAGE (5%, 59:1 acrylamide:bis-acrylamide ratio). The purified nucleosomes were stored in a buffer containing 10 mM Tris (pH-7.5) and 1 mM EDTA at 4 °C.

### Purification of recombinant UDG and HMCES

A pET-His-GFP vector (N-terminal His-GFP tag) with the human uracil DNA glycosylase (UDG) gene was obtained from GenScript. The pET-His-GFP-UDG vector transformed into BL21-CodonPlus (DE3)-RIPL *E. coli* cells (Agilent). The transformed cells were grown at 37 °C to an OD600 of 0.6 and UDG expression induced using 1.0 mM IPTG for 18 h at 18 °C. The cells were lysed via sonication in a buffer containing 25 mM HEPES (pH 7.5), 200 mM NaCl, 0.1 mM TCEP, 20 mM Imidazole, and a cocktail of protease inhibitors (AEBSF, leupeptin, benzamidine, pepstatin A). The lysate was clarified via centrifugation and loaded onto a HisTrap HP column (Cytiva) equilibrated with 25 mM HEPES (pH 7.5), 200 mM NaCl, 0.1 mM TCEP, and 20 mM Imidazole, and eluted in a buffer 25 mM HEPES (pH 7.5), 200 mM NaCl, 0.1 mM TCEP, and 400 mM Imidazole. The UDG protein was then liberates from the His-GFP tag via TEV protease and further purified using a HiPrep SP HP column (Cytiva) equilibrated with 25 mM HEPES (pH 7.5), 50 mM NaCl, and 0.1 mM TCEP, and eluted in a buffer 25 mM HEPES (pH 7.5), 1 M NaCl, and 0.1 mM TCEP. The resulting UDG protein was combined and purified via size exclusion chromatography using a HiPrep Sephacryl S-200 16/60 HR (Cytiva) in a buffer containing 20 mM HEPES (pH 7.5), 150 mM NaCl, and 0.1 mM TCEP. The purity of the UDG proteins were confirmed via denaturing SDS-PAGE and the purified UDG protein stored long term at −80 °C.

Recombinant human HMCES protein was purified using a previously described protocol^155^. In brief, a pNIC-HMCES vector was transformed in *E. coli* cells. Cells were grown in terrific broth (TB) medium at 37 °C to an OD_600_ of 0.7 and HMCES expression induced by the addition of 0.5 mM IPTG for 4 hrs. Cell pellets were collected via centrifugation and resuspended in a buffer containing 20 mM HEPES/KOH (pH 7.5), 500 mM KCl, 5 mM MgCl₂, 30 mM imidazole, 10% glycerol, 0.1% IGEPAL, 0.04 mg/ml Pefabloc SC, 1 mM TCEP and cOmplete EDTA-free protease inhibitor cocktail. The lysate was loaded onto a Strep-Tactin® XT Superflow® high-capacity cartridge, washed with a buffer containing 20 mM HEPES/KOH pH 7.5, 500 mM KCl, 5 mM MgCl₂, 1 mM TCEP, and eluted in a buffer containing 20 mM HEPES/KOH pH 7.5, 500 mM KCl, 5 mM MgCl₂, 1 mM TCEP, and 50 mM biotin. HMCES was further purified using a HiTrap Heparin HP columns and eluted with buffer containing 20 mM HEPES/KOH pH 7.5, 1 M KCl, 5 mM MgCl₂, 1 mM TCEP. The HMCES protein was combined and further purified using a HiPrep Sephacryl S-200 16/60 HR (Cytiva) in a buffer containing 20 mM HEPES/KOH pH 7.8, 150 mM KCl, 5 mM MgCl₂, 10% glycerol, 1 mM TCEP. The purity of the HMCES protein was confirmed via denaturing SDS-PAGE and purified HMCES protein stored long term at −80 °C.

### Generation of HEMCES-DPC and HEMCES-DPC nucleosome formation

AP-site DNA was generated via enzymatic digestion of dU-containing oligos (see Supplementary Table 1. The dU-containing oligonucleotides (10 µM) were incubated with recombinant UDG (10 µM) for 2 hours at 25 °C to excise the uracil and generate an AP-site. The oligonucleotides were then incubated with recombinant HMCES (20 µM) for 2 hours at 25 °C, and the HMCES-DPC stabilized via sodium borohydride (NaBH₄) crosslinking for an additional 2 hours at 25 °C. The HEMCES-DPC-DNA was loaded onto a HiPrep Q HP column (Cytiva) equilibrated with 50 mM HEPES (pH 7.5) and 400 mM NaCl, and eluted in a buffer 50 mM HEPES (pH 7.5) and 2 M NaCl. Fractions containing the HMCES-DPC-DNA were combined, exchanged into a buffer containing 20 mM Tris-HCl (pH 7.5), 1 mM EDTA, 1 M NaCl, and 1mM DTT, and stored short-term at 4 °C prior to nucleosome reconstitution.

To generate HMCES-DPC-DNA nucleosomes, HMCES-DNA, H2A/H2B dimer, and H3/H4 tetramer were mixed at a ratio of 1:2.2:1, respectively, in a buffer containing 20 mM Tris-HCl (pH 7.5), 1 M NaCl, 1 mM EDTA, and 1mM DTT. Nucleosomes were assembly via a stepwise decrease in NaCl concentration over a 24-hour period - from 1 M NaCl to 0.8 M NaCl, to 0.6 M NaCl, to 0.4 M NaCl, to 0.2 M NaCl, and to 0 M NaCl. The formation of the HMCES-DPC nucleosomes was assessed via native PAGE gels (5%, 59:1 acrylamide:bis-acrylamide ratio).

## FRET assay

### Oligonucleotides in FRET assay

Oligonucleotides comprising the single strand DNA (ssDNA) gap substrate (**Table 1**) were synthesized by Integrated DNA Technologies (Coralville, IA) or Bio-Synthesis (Lewisville, TX) and purified on denaturing polyacrylamide gels. The concentrations of unlabeled DNAs were determined from the absorbance at 260 nm using the provided extinction coefficients. Concentrations of Cyanine-labeled DNAs were determined from the extinction coefficient of their respective label (Cy3 at 550 nm, ε_550_ = 150,000 M^−1^cm^−1^, Cy5 at 650 nM, ε_650_ = 250,000 M^−1^cm^−1^). For annealing the ssDNA gap substrate, the template strand and the cyanine-labeled primer strands were mixed in equimolar amounts in 1X annealing buffer (10 mM TrisHCl, pH 8.0, 100 mM NaCl, 1 mM EDTA), heated to 95 °C for 5 minutes, and allowed to slowly cool to room temperature.

**Table 1.**
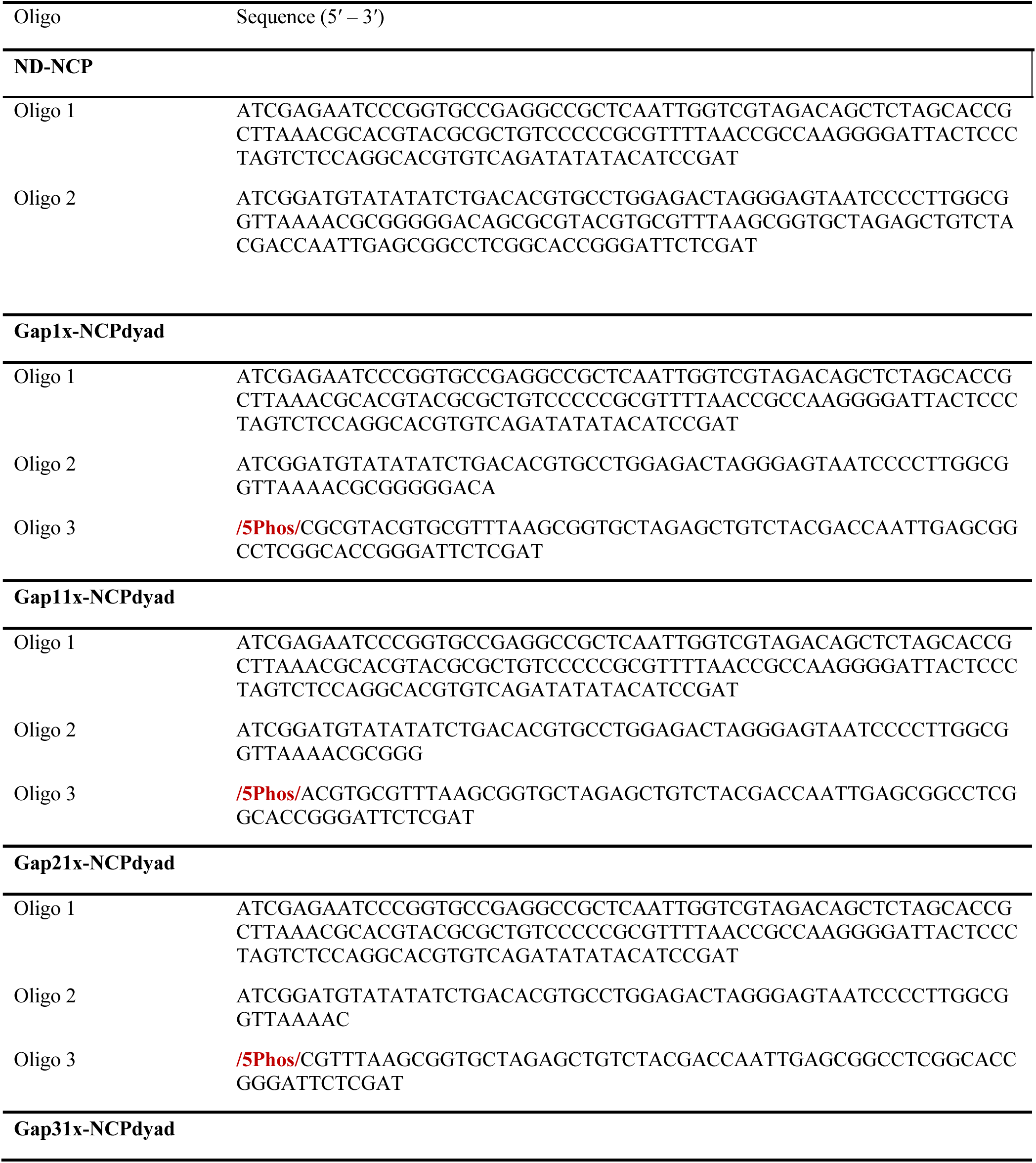
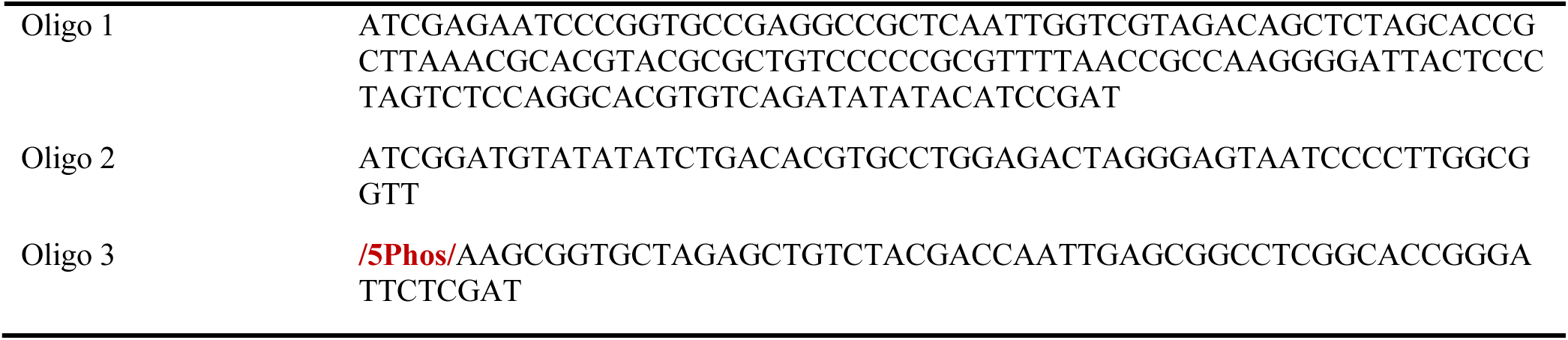
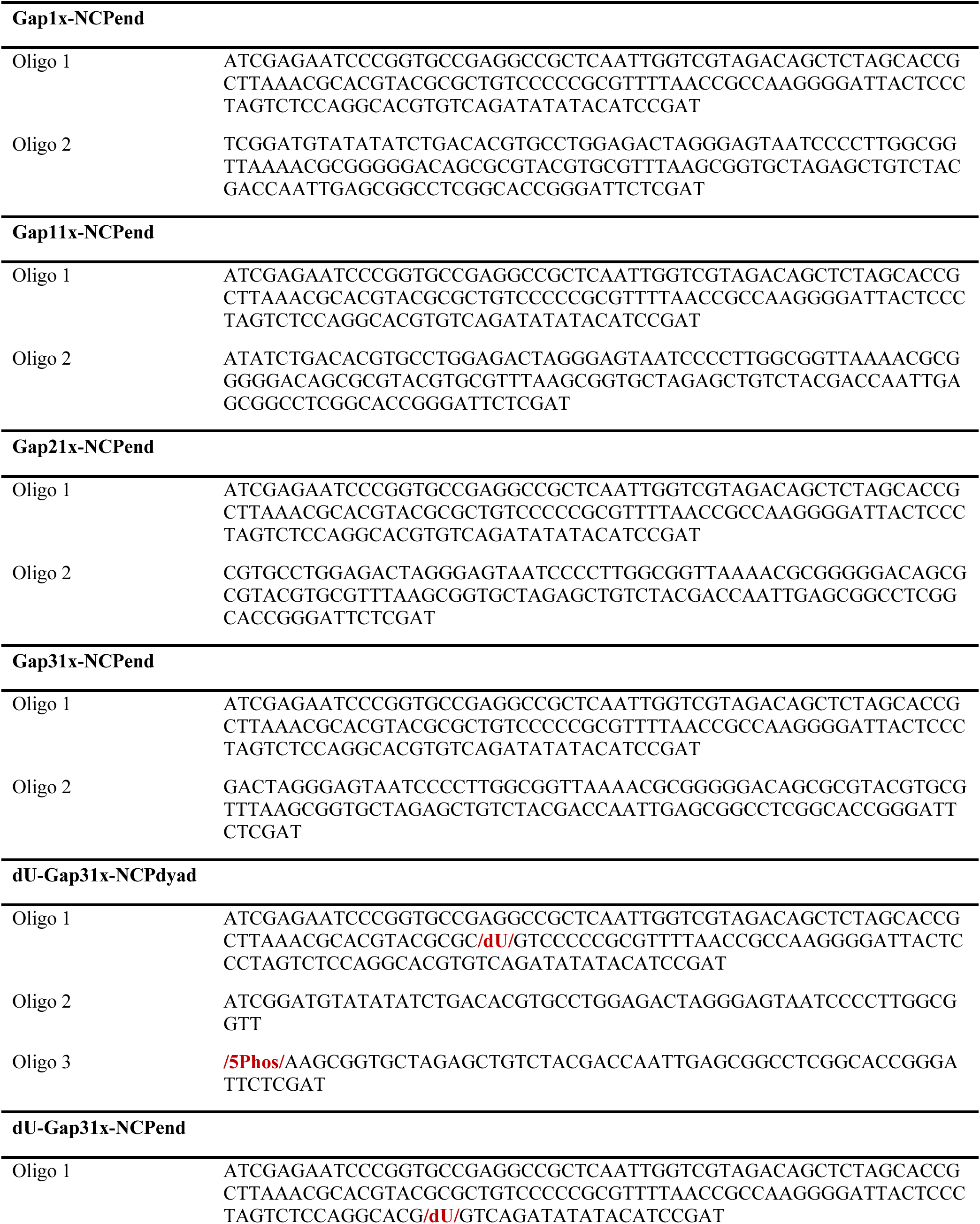
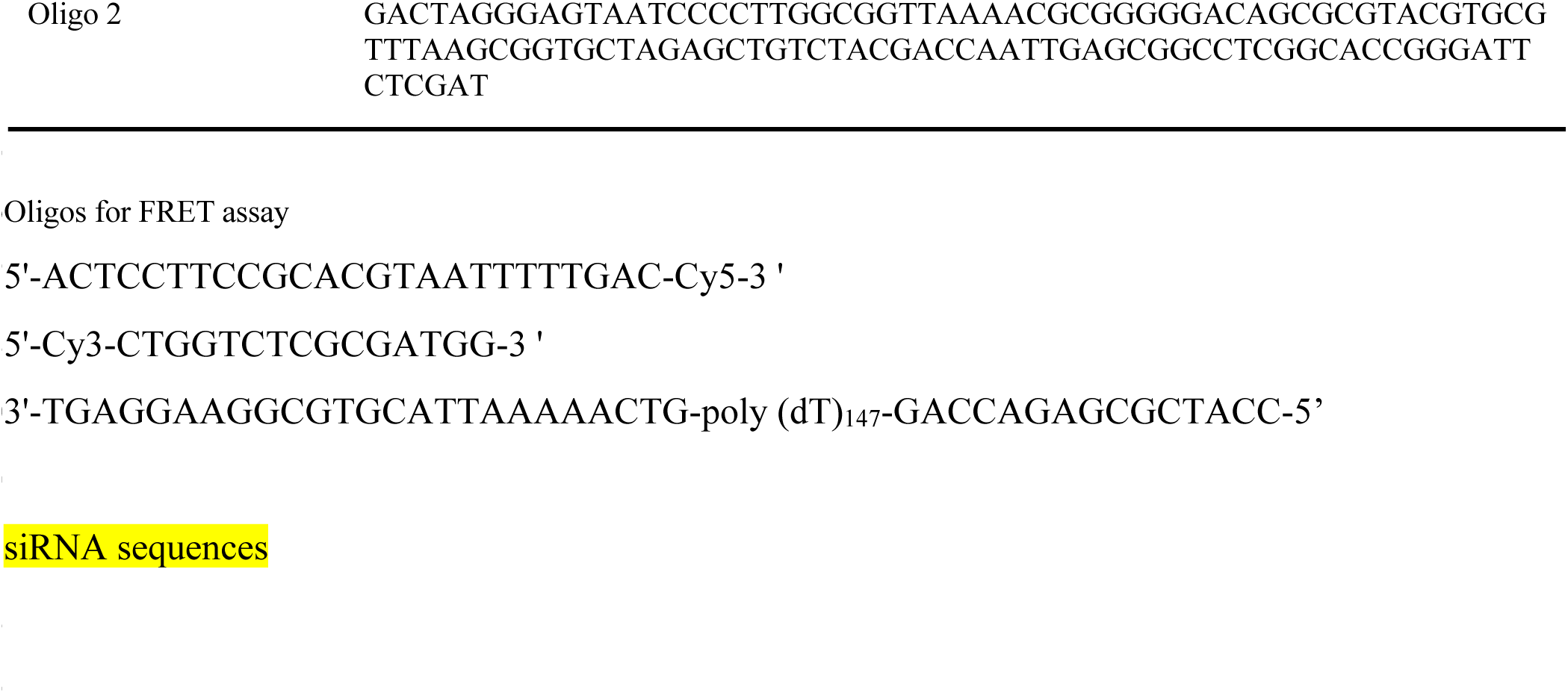
Oligo sequences in this study.

### Recombinant Human Proteins in FRET assay

Human RPA was obtained as previously described ^156^. The concentration of active RPA was determined via a FRET-based activity assay as described previously ^109^. Human recombinant histone H3.1 – H4 tetramers (Cat. # 14-1056) and histone H2A-H2B dimers (Cat. # 10-1052) were purchased from EMD Millipore (Temecula, CA) and utilized to reconstitute nucleosomes on the ssDNA gap substrate via stepwise salt dilution dialysis method, as described previously ^157^. Assembled nucleosomes were purified and verified by native PAGE, as described previously ^157^.

### FRET Experiments

All experiments were performed in a 16.100F-Q-10/Z15 sub-micro fluorometer cell (Starna Cells) at room temperature (23 ± 2 °C) in 1X Replication Buffer (25 mM TrisHCl, pH 7.5, 125 mM KOAc, 10 mM Mg(OAc)_2_) supplemented with 1 mM DTT, 0.1 mg/mL BSA, and the ionic strength adjusted to physiological (200 mM) by addition of KOAc. Fluorescence intensities were monitored in a Horiba Scientific Duetta-Bio fluorescence/absorbance spectrometer with excitation and emission slit widths set at 10 nm. For all experiments described below the final concentrations of all reaction components are indicated. The concentration of RPA (6.76 μM) utilized in the experiments described below agrees with the concentration of RPA in the nucleoplasm of a human cell (10 μM range)^103–105^.

For steady state (equilibrium) FRET assays in Figure S5B, a DNA substrate (100 nM ssDNA Gap or ssDNA Gap Nuc) is pre-incubated for 5 minutes in the absence or presence of RPA (6.76 μM heterotrimer). Under these conditions, the concentration of RPA is ∼11-fold higher than the concentration of RPA-binding sites within the the ssDNA Gap DNA substrate. The solution is then transferred to a fluorometer cell and the cell is placed in the instrument. Next, the solution is excited at 514 nm and the fluorescence emission intensities (*I*) are monitored essentially simultaneously (Δt = 0.118 ms) at the peak emission wavelengths for Cy3 (563 nm, *I*_563_) and Cy5 (665 nm, *I*_665_) over time until both stabilize for at least 1 min. The *I*_665_ and *I_563_* values within this stable region are utilized to calculate the approximate FRET efficiencies (*E*_FRET_) from the equation 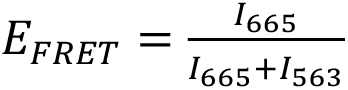 and the resultant values are averaged to obtain the *E*_FRET_ value observed for a given condition at equilibrium. Finally, the solution is excited at 514 nM and the fluorescence emission spectra is recorded from 530 to 750 nm.

For pre-steady state FRET assays (Figure S5C) ssDNA Gap Nuc (100 nM) is transferred to a fluorometer cell and the cell is placed in the instrument. The solution is excited at 514 nm and *I*_563_ and *I*_665_ are monitored over time (as described above) until both stabilize for at least 1 min. The *I*_665_ and *I_563_* values within this stable region are utilized to *E*_FRET_ and the resultant values are averaged to obtain the *E*_FRET_ value observed prior to the addition of RPA. Next, RPA (6.76 μM heterotrimer) is added, the resultant solution is mixed, and *I*_665_ and *I_563_* are monitored beginning 30 s after the addition of RPA. All recorded fluorescence emission intensities are corrected by a respective dilution factor. For all recordings of the fluorescence emission intensities (*I_665_* and *I_563_*), *E*_FRET_ values are calculated as described above and plotted as a function of time after RPA is added.

For all plots in all figures, each data point/column/spectra represent the average ± standard error of the mean (SEM). Error bars are present for all data points on all plots in all figures but may be smaller than the data point.

### Quantification and statistical analysis

The statistical test was performed using GraphPad Prism and R software. The statistics for the high-content microscopy experiments was performed on the mean of individual replicates. Statistical analyses were completed using Prism. A Kruskal-Wallis test was used for experiments with more than two samples, and P values were calculated by Prism for the multiple comparisons. A two-tailed t test was used to compare two samples with normally distributed data. No statistical methods or criteria were used to estimate sample size or to include/exclude samples. Statistical details of individual experiments can be found in the figure legends and in Results. Experiments shown are representative of at least two biological replicates unless otherwise indicated in the figure legend.

## Acknowledgement

We would like to thank David Cortez Lab for the SMARCAL1, HLTF, ZRANB3 single and triple knockout U2OS cell lines; Alessandro Vindigni lab for the pcDNA3.1/nV5-DEST PrimPol WT and Mutants vectors; Weixing Zhao lab for the MDA-MB-436 and BRCA1 complemental cell lines, DLD-1 WT and BRCA2 knockout cell lines. Part of this work was supported by funding to M. H. from the National Institutes of Health to M.H. (R35 GM147238-03).

## Author contribution

W.L. conceptualized the project. Q.W. and C.Z. performed PLA, IF, iPOND, western blot assays.

C.Z. performed data analysis for genomic and proteomic experiments. Y.D. and T.Z. performed Mass spectrometry experiments. N.I.M. and T.M.W. performed in vitro nucleosome assembly assays. M.G. and S.T. purified HMCES proteins and performed RADAR assay. T.H.L. and M.H. performed the FRET assay. W.L., Q.W., T.M.W., and M.H. wrote the manuscript, T.M.W and TP.Z. proofread the manuscript with contributions from all the authors.

## Supplementary Figure Legends

**Figure S1.**
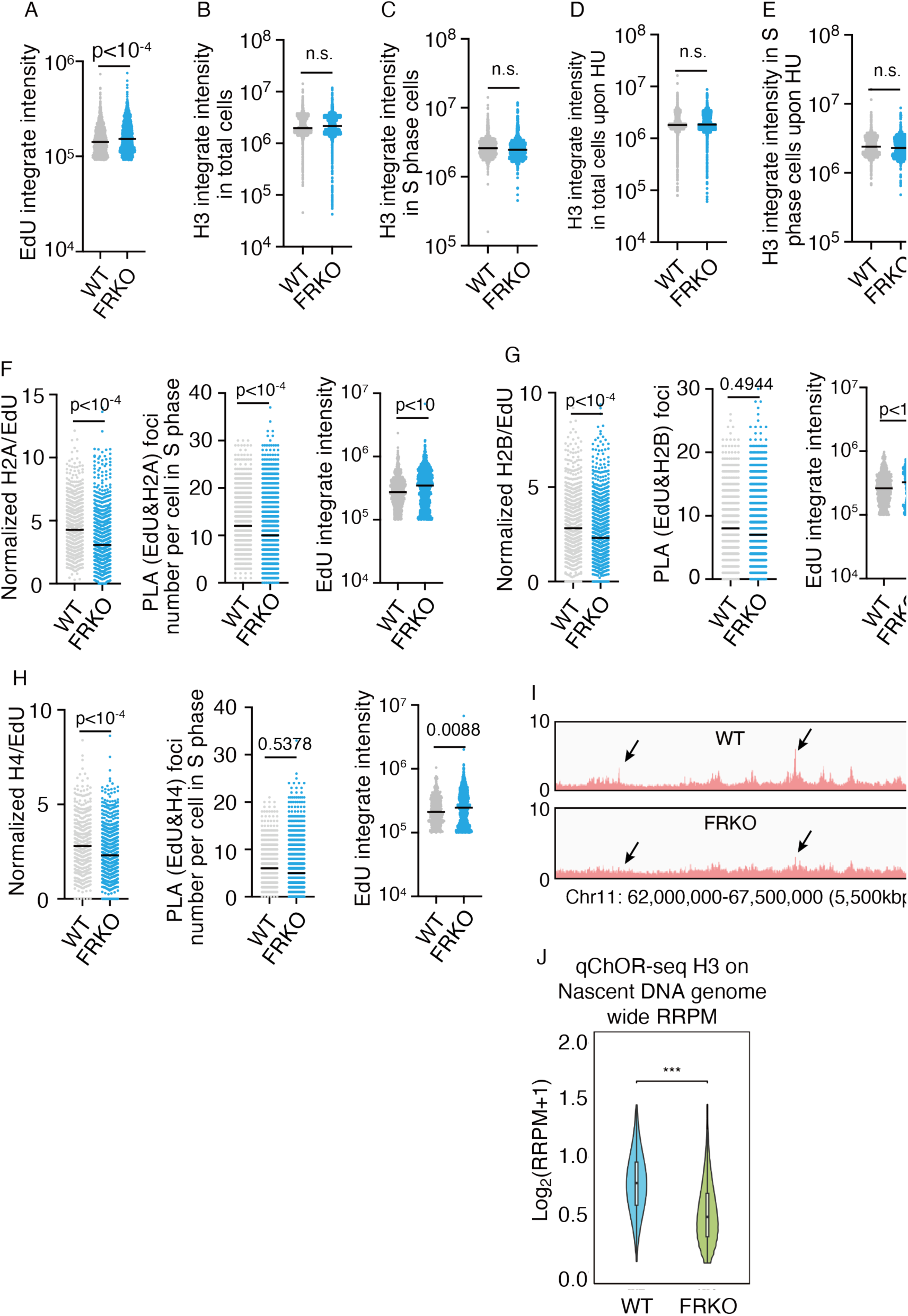
Fork reversal deficiency reduces nucleosome density on nascent DNA (related to Figure 1). FRKO clone #29 is used in figure S1A-J. A) EdU incorporation in WT and FRKO cells treated with EdU + HU (as in Fig. 1B). EdU detected by azide-488 click reaction. B–E) Immunofluorescence for H3 in WT and FRKO cells under indicated conditions. N = 3; p-values from two-tailed unpaired t-test. F–H) PLA for EdU and H2A (G), H2B (H), and H4 (I). Left: PLA foci/cell; middle: EdU intensity; right: normalized PLA/EdU. ≥1,000 cells/sample. N ≥ 3. I) Genome browser views of H3 ChOR-seq from the same experiment in Figure 1E, normalized to reads per million using exogenous spike-in. N = 2. J) Quantitative ChOR-seq for H3. Average profiles across H3 peaks in WT and FRKO cells, normalized by reference-adjusted RPM using D. melanogaster spike-in. N = 2; Wilcoxon signed-rank test.

**Figure S2.**
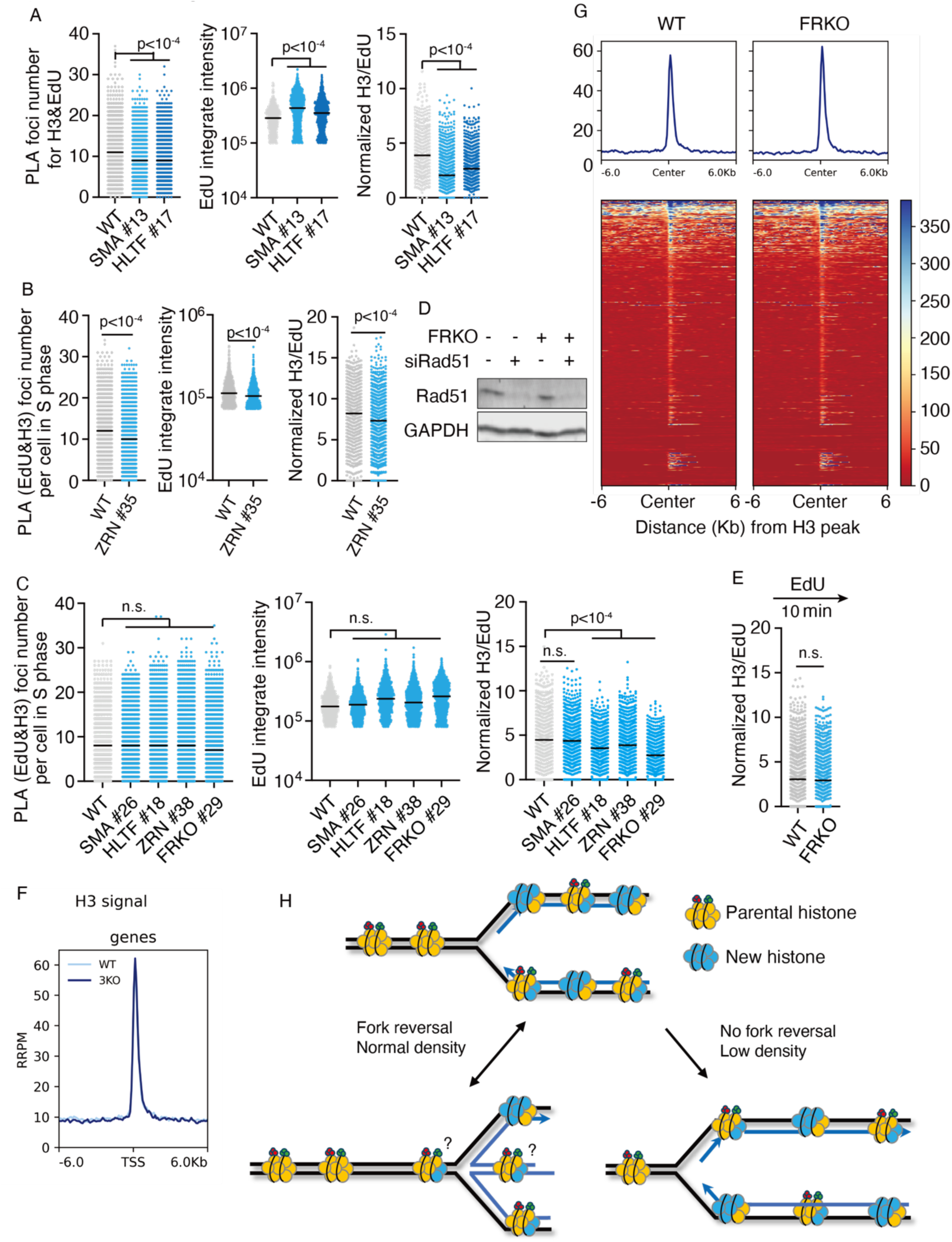
Low nucleosome density phenotype in individual fork reversal enzymes knockout clones of U2OS cell lines (related to Figure 1). A–C) PLA for EdU and H3 in individual fork reversal enzyme KO clones. N = 3. p-values from unpaired t-test in B) and Kruskal–Wallis test in A) and C). D) Western blot showing siRAD51 knockdown efficiency. E) PLA for EdU and H3 in WT and FRKO cells without HU. Cells labeled with EdU (10 min) and harvested for PLA. N = 3; p-values from unpaired t-test. F) ChOR-seq analysis of H3 occupancy in WT and FRKO cells without HU, normalized to spike-in chromatin. The cells are treated same as in D). G) Heatmaps of H3 ChOR-seq signal (same data from F) in unperturbed condition. N = 2. G) Model schematic: fork reversal prevents low nucleosome density under replication stress.

**Figure S3.**
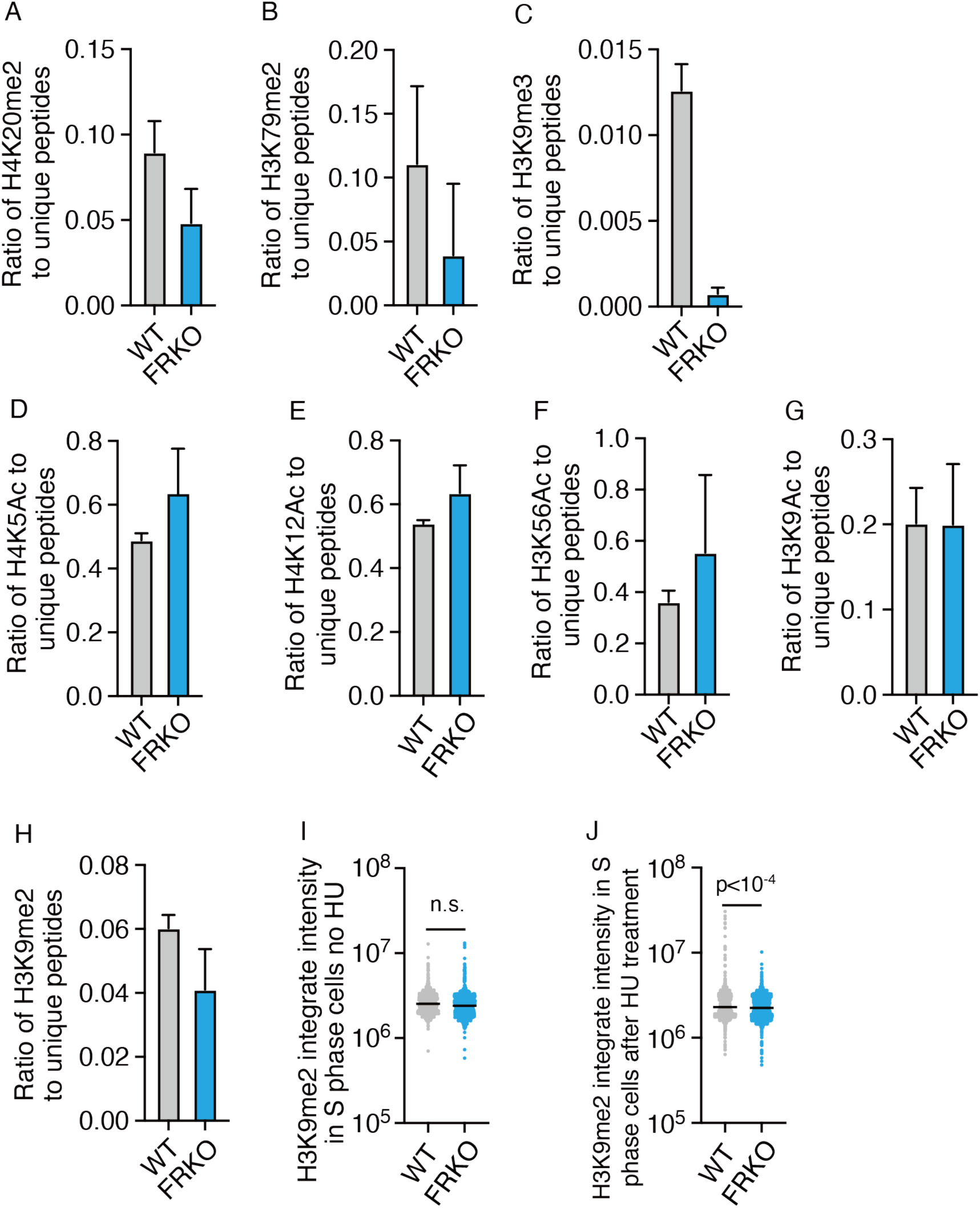
Fork reversal deficiency leads to parental histone loss (related to Figure 2). A-H) Histone modifications quantified in WT and FRKO cells by iPOND–Propionylation-MS. Modification level equals modified peptide count/unique peptide count. N = 3; error bars is generated from standard deviation. I-J) Immunofluorescence for H3K9me2 in indicated conditions. N = 3; unpaired t-test.

**Figure S4.**
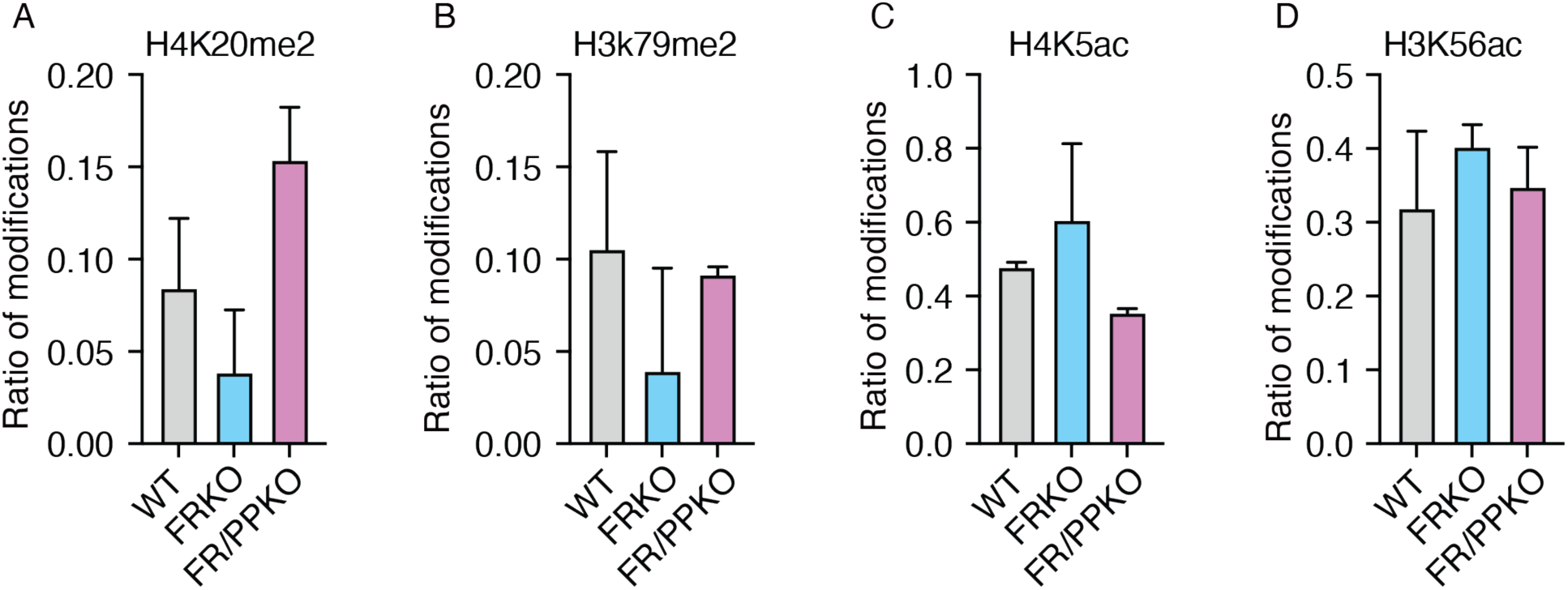
PrimPol depletion rescues parental histone loss and new histone increase, related to Figure 3. A-D) Histone modifications quantified in WT, FRKO, and FR/PPKO cells by iPOND–Propionylation-MS. Modification level equals modified peptide count/unique peptide count. N = 3; error bars is generated from standard deviation.

**Figure S5.**
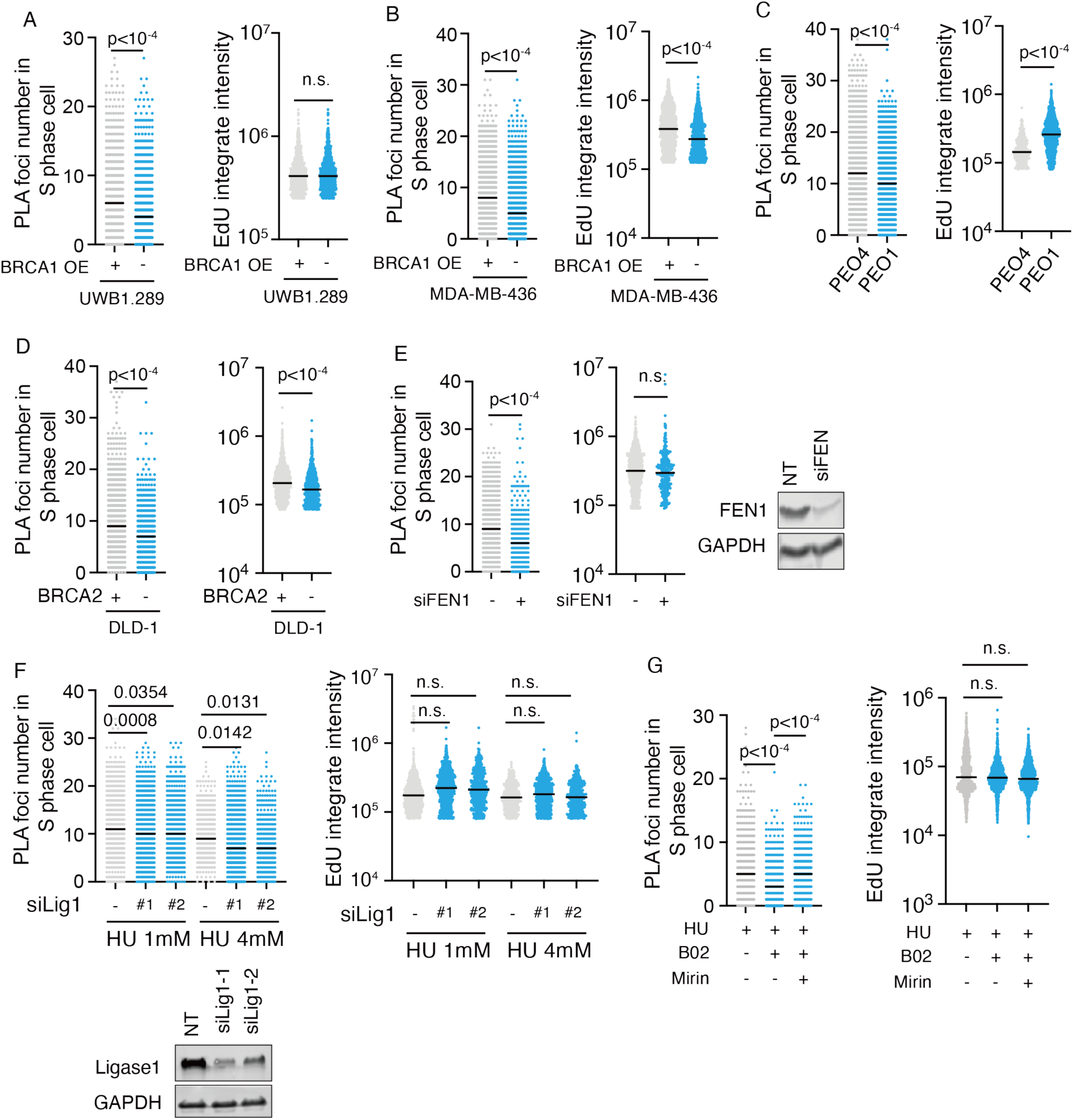
ssDNA gaps loosen the nucleosome assembly on nascent DNA in cells (related to Figure 4). A–G) PLA for EdU and H3 in indicated treatments. PLA foci number for each cell and EdU intensity as shown (from the same data as Fig. 4). N = 3; p-values from unpaired t-test.

**Figure S6.**
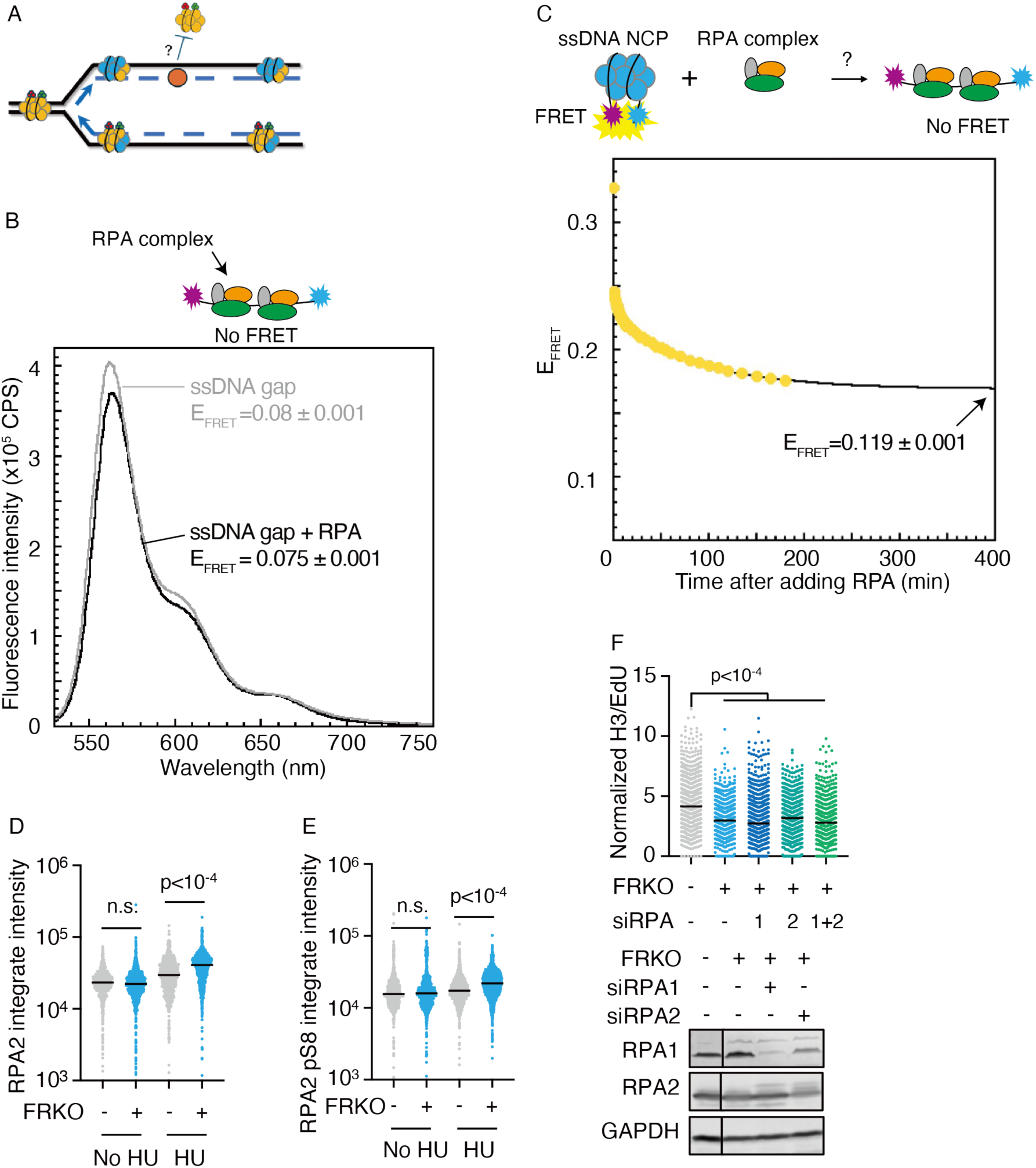
RPA evicts nucleosome from ssDNA in vitro but doesn’t not rescue the low nucleosome density in cells. A) Model schematic. B) FRET signal for ssDNA gap and RPA bound ssDNA gap. Saturating the ssDNA within the ssDNA Gap DNA substrate with RPA does not significantly affect the *E*_FRET_ observed at equilibrium. Fluorescence emission spectra obtained by excitation at 514 nm is shown. Each spectra is the average of at least two independent spectra with the SEM shown in grey. The average *E*_FRET_ values observed at equilibrium for each condition are shown in the inset. Each reported value is the average of at least 4 independent measurements (±SEM). C) Schematic representation of the FRET pair and assay to monitor unwrapping of ssDNA from around histone octamers within ssDNA Gap nucleosomes. In ssDNA Gap nucleosomes conditions as described in Figure 5E, RPA is added and E_FRET_ values are monitored over time. Trace is the mean of at least three independent traces. D-E) Immunofluorescence for RPA2 and phosphorylation on RPA2-S8 in WT and FRKO U2OS cells. The cells are treated same as in Figure 6A. N = 3. F) PLA for EdU and H3 in siRPA1 and/or siRPA2 cells (treated as in Fig. 1B). N ≥ 5; Kruskal–Wallis test. Western blot showed siRNA knockdown efficiency.

**Figure S7.**
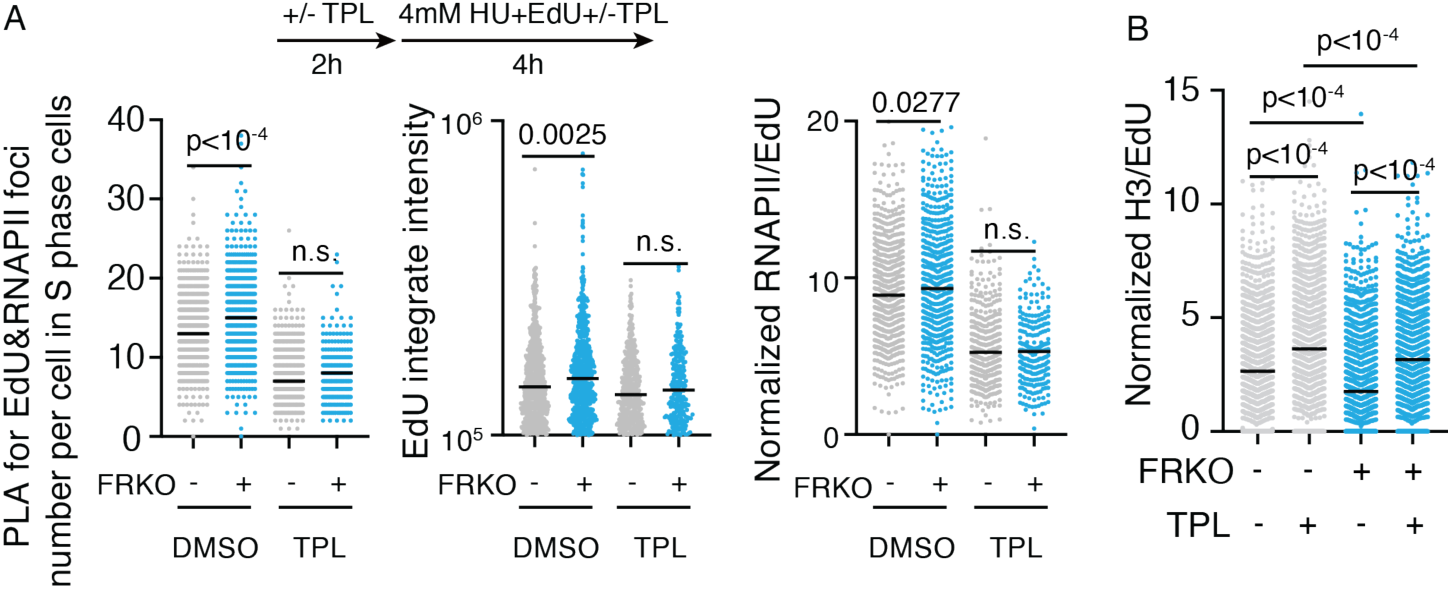
Transcription reduces nucleosome density in both WT and FRKO cells. A) PLA for EdU and RBP1 in indicated treatments. Left, PLA foci number for each cell and EdU intensity as shown. Middle, EdU integrity intensity for each cell in the indicated treatment. Right, normalized PLA signal in each cell in the indicated treatment. TPL concentration is 10μM. N = 3; p-values from Kruskal-Wallis test. B) PLA for EdU and H3 in indicated treatments. N = 3; TPL concentration is 10μM. p-values from Kruskal–Wallis test.

**Figure S8.**
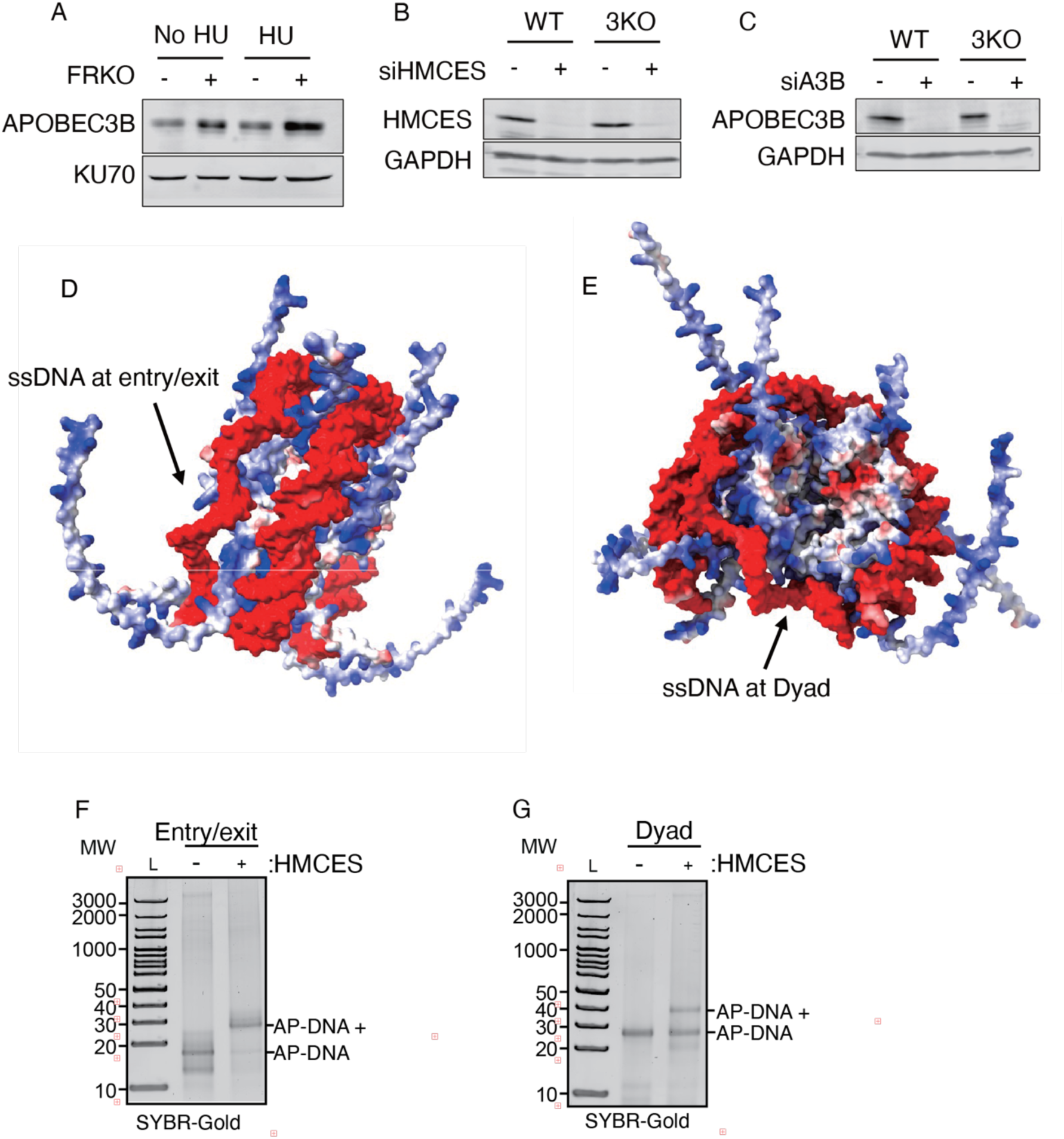
HMCES-DPC at ssDNA gaps reduces nucleosome density in FRKO cells (related to Figure 7). A) Western blot for chromatin-bound APOBEC3B in WT and FRKO cells with or without HU treatment. B-C) Western blot for HMCES (B) and APOBEC3B (C) to confirm knockdown efficiency in WT and FRKO cells. D-E) Structural models of ssDNA-gapped nucleosomes predicted by AlphaFold. D) ssDNA gap at entry/exit. E) ssDNA gap at dyad. F-G) HMCES-ssDNA crosslink preparation. Gels show HMCES–DNA crosslink products. AP-DNA represents DNA that contains abasic sites; L represents ladder.

